# HOXA9 promotes MYC-mediated leukemogenesis by maintaining gene expression for multiple anti-apoptotic pathways

**DOI:** 10.1101/2020.10.20.347765

**Authors:** Ryo Miyamoto, Akinori Kanai, Hiroshi Okuda, Satoshi Takahashi, Hirotaka Matsui, Toshiya Inaba, Akihiko Yokoyama

## Abstract

HOXA9 is often highly expressed in leukemias. However, its precise roles in leukemogenesis remain elusive. Here, we show that HOXA9 maintains gene expression for multiple anti-apoptotic pathways to promote leukemogenesis. In *MLL*-rearranged leukemia, MLL fusion directly activates the expression of MYC and HOXA9. Combined expression of MYC and HOXA9 induced leukemia, whereas single gene transduction of either did not, indicating a synergy between MYC and HOXA9. HOXA9 sustained expression of the genes implicated to the hematopoietic precursor identity when expressed in hematopoietic precursors, but did not reactivate it once silenced. Among the HOXA9 target genes, *BCL2* and *SOX4* synergistically induced leukemia with *MYC*. Not only BCL2, but also SOX4 suppressed apoptosis, indicating that multiple anti-apoptotic pathways underlie cooperative leukemogenesis by HOXA9 and MYC. These results demonstrate that HOXA9 is a key transcriptional maintenance factor which promotes MYC-mediated leukemogenesis, potentially explaining why HOXA9 is highly expressed in many leukemias.

## Introduction

Mutations of transcriptional regulators often cause aberrant gene regulation of hematopoietic cells, which leads to leukemia. Structural alterations of the mixed lineage leukemia gene (*MLL* also known as *KMT2A*) by chromosomal translocations cause malignant leukemia that often associates with poor prognosis despite the current intensive treatment regimens (1). MLL encodes a transcriptional regulator that maintains segment-specific expression of homeobox (*HOX*) genes during embryogenesis (2), which determines the positional identity within the body (3–5). During hematopoiesis, MLL also maintains the expression of posterior *HOXA* genes and *MEIS1* (another homeobox gene), which promote the expansion of hematopoietic stem cells and immature progenitors (6–10). The oncogenic MLL fusion protein constitutively activates its target genes by constitutively recruiting transcription initiation/elongation factors thereto (11–13). Consequently, *HOXA9* and *MEIS1* are highly transcribed in *MLL*-rearranged leukemia (7). Forced expression of HOXA9 (but not MEIS1) immortalize hematopoietic progenitor cells (HPCs) ex vivo (14, 15). Co-expression of HOXA9 with MEIS1 causes leukemia in mice which recapitulates *MLL*-rearranged leukemia (15). Moreover, HOXA9 is highly expressed in many non-*MLL*-rearranged leukemias such as those with *NPM1* mutation and *NUP98* fusion and is associated with poor prognosis (16). These findings highlight HOXA9 as a major contributing factor in leukemogenesis. Nevertheless, the mechanism by which HOXA9 promotes oncogenesis remains elusive.

HOXA9 is considered to function as transcription factor, which retains a sequence-specific DNA binding ability. HOX proteins have an evolutionally conserved homeodomain which possesses strong sequence preferences (17). HOXA9 associates with other homeodomain proteins such as PBX and MEIS family proteins (14, 18). HOXA9 and those HOXA9 cofactors form a stable complex on a DNA fragment harboring consensus sequences for each homeodomain protein (18, 19), suggesting that they form a complex of different combinations in a locus-specific manner depending on the availability of the binding sites. Recently, it has been reported that HOXA9 specifically associates with enhancer apparatuses (e.g. MLL3/4) to regulate gene expression (20–22). However, the mechanisms by which HOXA9 activate gene expression remain largely unclear.

In this study, we reveal the oncogenic roles for HOXA9 and its target gene products in leukemogenesis and its unique mode of function as a transcriptional maintenance factor which preserves an identity of a hematopoietic precursor.

## Results

### MLL fusion proteins and HOXA9 sustain MYC expression against differentiation-induced transcriptional suppression

To identify the direct target genes of MLL fusion proteins, we first examined the genome-wide localization pattern of MLL fusion proteins by chromatin immunoprecipitation (ChIP) followed by deep sequencing (ChIP-seq), using HB1119 cells, a cell line expressing a fusion gene of *MLL* and *ENL* (*MLL-ENL)*. We observed MLL ChIP signals on the *MYC*, *HOXA9*, *HOXA10*, and *MEIS1* loci (Figure 1A) (23), which were further confirmed by ChIP-qPCR analysis (Supplemental Figure 1).

**Figure 1.**
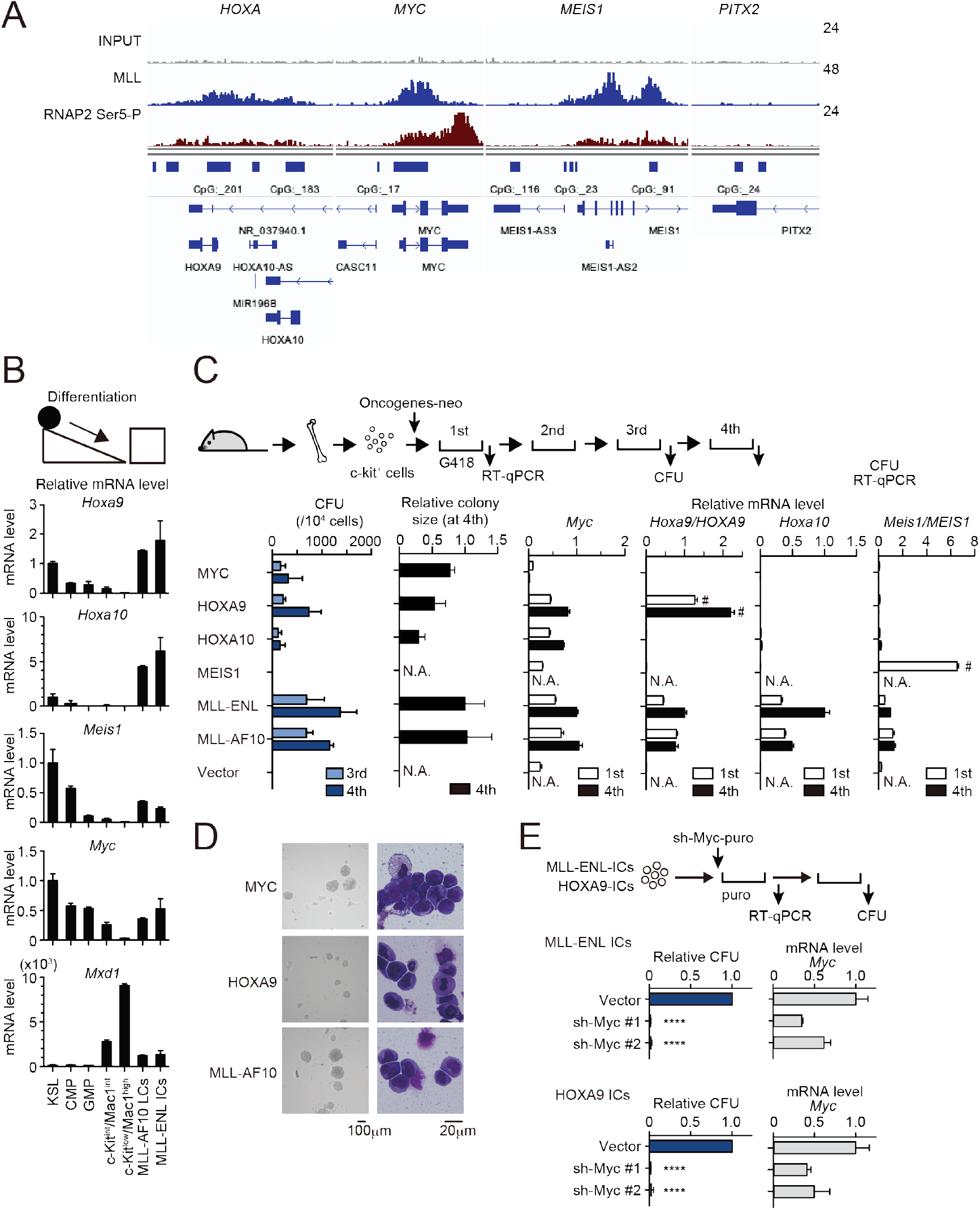
MLL fusion proteins and HOXA9 sustain MYC expression against differentiation-induced transcriptional suppression. **A**, Genomic localization of MLL-ENL in HB1119 cells. ChIP signals at the loci of posterior *HOXA* genes, *MEIS1*, *MYC*, and *PITX2* (negative control) are shown using the Integrative Genomics Viewer (The Broad Institute). **B**, Expression of MLL-target genes and *Mxd1* (a differentiation marker) during myeloid differentiation. Bone marrow cells at various differentiation stages were obtained by FACS sorting and analyzed by RT-qPCR. Expression levels relative to KSL are shown (Mean with SD, n = 3, PCR replicates). MLL-AF10-LCs and MLL-ENL-ICs were included in the analysis for comparison. KSL: c-Kit^+^, Sca1^+^, and Lineage^−^; CMP: common myeloid progenitor; GMP: granulocyte macrophage progenitor; int: intermediate. **C**, Transforming potential of MLL target genes. Clonogenic potential of the indicated constructs was analyzed by myeloid progenitor transformation assays. Colony forming unit per 10^4^ cells (CFU) (Mean with SD, n = 3, biological replicates), relative colony size (Mean with SD, n ≥ 100), and relative mRNA levels of indicated genes (Mean with SD, n = 3, PCR replicates) were measured at the indicated time points. #: Both endogenous murine transcripts and exogenous human transcripts were detected by the qPCR primer set used. N.A.: not assessed. **D**, Morphologies of the colonies and transformed cells. Representative images of bright field (left) and May-Grunwald-Giemsa staining (right) are shown with scale bars. **E**, Effects of *Myc* knockdown on MLL-ENL- and HOXA9-ICs. Relative CFU (Mean with SD, n = 3, biological replicates) and mRNA level of *Myc* (Mean with SD, n = 3 PCR replicates) are shown. Statistical analysis was performed using ordinary one-way ANOVA with the vector control. \*\*\*\**P* < 0.0001.

To characterize the dynamic changes in MLL target gene expression during differentiation, we isolated bone marrow cells from mice at various differentiation stages ranging from the most immature population (c-Kit^+^, Sca1^+^, Lineage^−^; KSL) containing hematopoietic stem cells to highly differentiated hematopoietic cells (c-Kit^low^/Mac1^high^) by fluorescence activated cell sorting (FACS) and performed qRT-PCR analysis. For comparison, we analyzed leukemia cells (LCs) harvested from mice suffering from MLL-AF10-induced leukemia (MLL-AF10-LCs) and immortalized cells (ICs) transformed by MLL-ENL ex vivo (MLL-ENL-ICs). In accordance with previous reports (7, 24), *Hoxa9*, *Hoxa10*, and *Meis1* were downregulated during the course of normal hematopoietic differentiation but remained abundant in MLL fusion-expressing cells (Figure 1B). *Myc* was highly expressed in KSL, common myeloid progenitors (CMP), and granulocyte/macrophage progenitors (GMP), which contain actively dividing populations (25), but was completely suppressed at highly differentiated c-kit^low^/Mac1^high^ stages. In the two MLL fusion-expressing cell lines, *Myc* was expressed at comparable levels to those in the progenitor fractions (CMP and GMP; Figure 1B). *Mxd1*, a differentiation marker, was highly expressed only in differentiated populations. These results indicate that *Myc*, *Hoxa9*, *Hoxa10*, and *Meis1* are intrinsically programmed to be silenced in normal hematopoietic differentiation but are aberrantly maintained by MLL fusion proteins.

To assess the oncogenic potential of MLL target genes, we performed myeloid progenitor transformation assays, wherein HPCs were isolated from mice, retrovirally transduced with each MLL target gene, and cultured in semi-solid medium supplemented with cytokines promoting myeloid lineage differentiation (Figure 1C, D). HPCs transduced with *MLL-ENL* and *MLL-AF10* produced a large number of colonies in the third and fourth passages with high mRNA levels of *Myc*, *Hoxa9*, *Hoxa10*, and *Meis1* in colonies of the first passage, confirming their potent transforming capacities. These cells were considered “immortalized” as they proliferate indefinitely in this ex vivo culture (26, 27). Ectopic expression of MYC immortalized HPCs; therefore, MYC expression is sufficient to induce proliferation of HPCs. Endogenous *Myc* expression was completely diminished in MYC-ICs, demonstrating that *Myc* expression is programed to be silenced and cannot be sustained by MYC itself. Ectopic expression of HOXA9 and HOXA10 (but not MEIS1) immortalized HPCs. Endogenous *Myc* expression was maintained in HOXA9/A10-ICs, indicating that HOXA9/A10 can sustain *Myc* expression against differentiation-induced transcriptional suppression. It should be noted that the colony size of HOXA9/A10-ICs was relatively small compared to MYC- or MLL fusion-ICs, suggesting a weaker proliferative potential. Knockdown of *Myc* by shRNA completely repressed the colony-forming ability of MLL-ENL-ICs and HOXA9-ICs (Figure 1E). Taken together, these results demonstrate that MLL fusion proteins and HOXA9 maintain *Myc* expression in spite of the differentiation-induced transcriptional suppression, and the maintenance of MYC expression is indispensable for immortalization of HPCs *ex vivo*.

### HOXA9 confers the identity of a hematopoietic precursor while MYC drives anabolic pathways

To identify the genes specifically regulated by HOXA9 but not by MYC, we performed RNA-seq analysis of HOXA9-ICs and MYC-ICs which do not express *HOXA9* (Figure 1C). Genes highly expressed in HOXA9-ICs but lowly expressed in MYC-ICs (defined as “HOXA9 high signature”) were associated with hematopoietic identity/functions, whereas genes highly expressed in MYC-ICs and lowly expressed in HOXA9-ICs (defined as “MYC high signature”) were associated with anabolic pathways (Figure 2A, B). MLL-AF10-ICs, which express endogenous *Hoxa9* and *Myc* at high levels (Figure 1C), expressed both HOXA9 high and MYC high signature genes (Figure 2A-C), indicating that MLL-AF10-ICs possess MYC-mediated highly proliferative potential and HOXA9-mediated hematopoietic identity. These differences between the HOXA9 high and MYC high signatures indicate that HOXA9 maintains the identity of a hematopoietic precursor.

**Figure 2.**
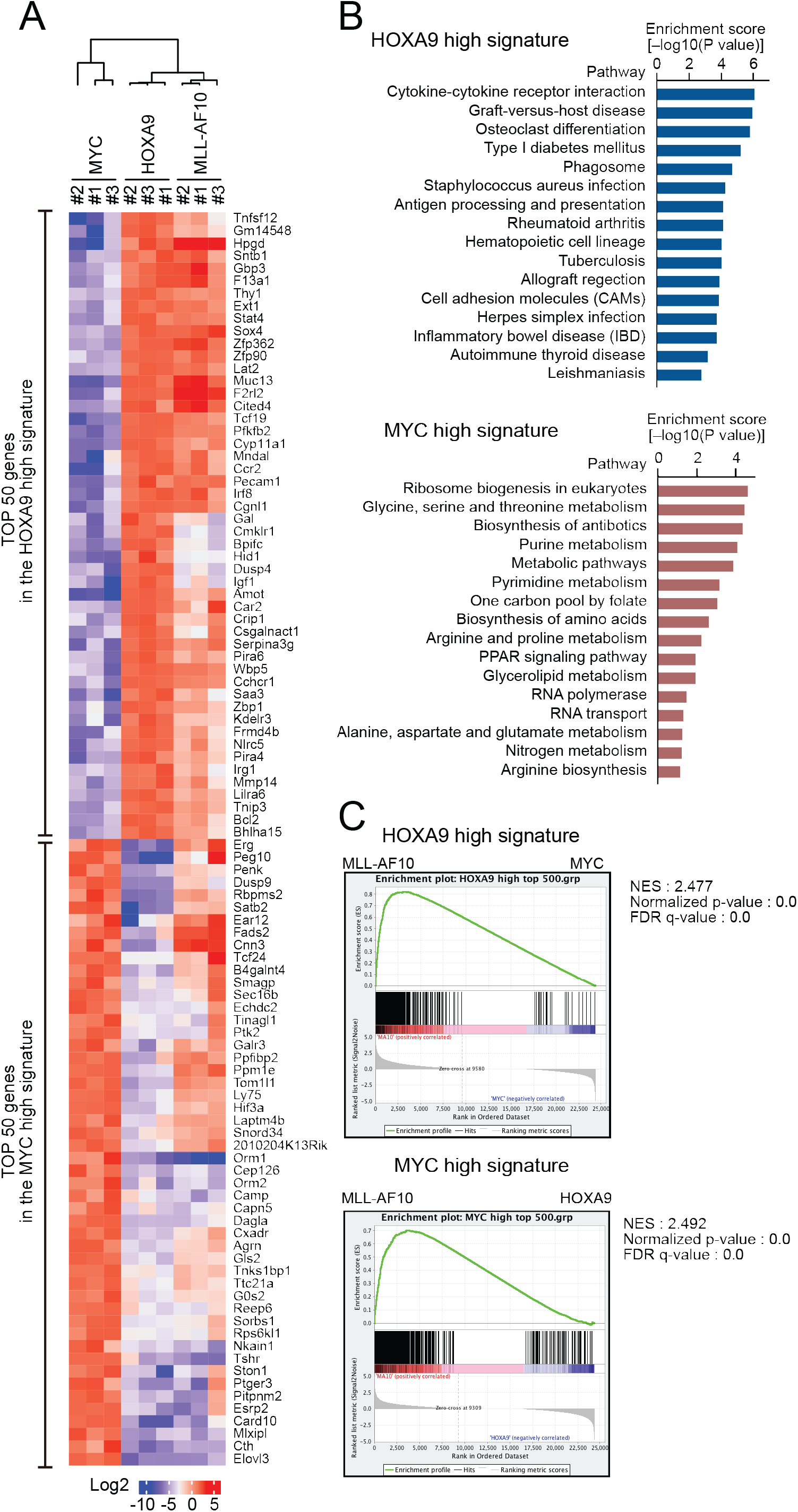
HOXA9 confers the identity of a hematopoietic precursor while MYC drives anabolic pathways. **A**, Relative expression of the top 50 genes categorized as the HOXA9 high signature and the MYC high signature in HOXA9-, MYC-, and MLL-AF10-ICs. **B**, Pathways related to the HOXA9 high signature (blue) and the MYC high signature (red). The top 500 genes in the HOXA9 high or MYC high signatures were subjected to KEGG pathway analysis. **C**, Gene set enrichment analysis of the HOXA9 high and MYC high signatures in MLL-AF10-ICs compared with MYC-ICs (top) and HOXA9-ICs (bottom), respectively.

### Apoptosis is induced by MYC, while is alleviated by HOXA9 and MLL-AF10

Given that excessive MYC activity promotes apoptosis (28), we evaluated their apoptotic tendencies in immortalized HPCs. MYC-ICs exhibited increased γH2AX, cleaved poly (ADP) ribose polymerase (PARP), and cleaved caspase 3 levels, indicative of a high degree of replication stress and apoptosis (Figure 3A). FACS analysis with Annexin V also showed that MYC-ICs had a larger apoptotic fraction than that of HOXA9-ICs (Figure 3B). MLL-AF10-ICs exhibited weak apoptotic tendencies similarly to HOXA9-ICs although their MYC expression tends to be higher than HOXA9-ICs (Figures 1C and 3A). Furthermore, most MYC-ICs underwent massive apoptosis 1d after cytokine removal, whereas HOXA9-ICs and MLL-AF10-ICs showed relative resistance (Figure 3C) and exhibited successful recovery of the live cell population after cytokine reintroduction (Supplemental Figure 2). These results indicate that HOXA9 confers anti-apoptotic properties.

**Figure 3.**
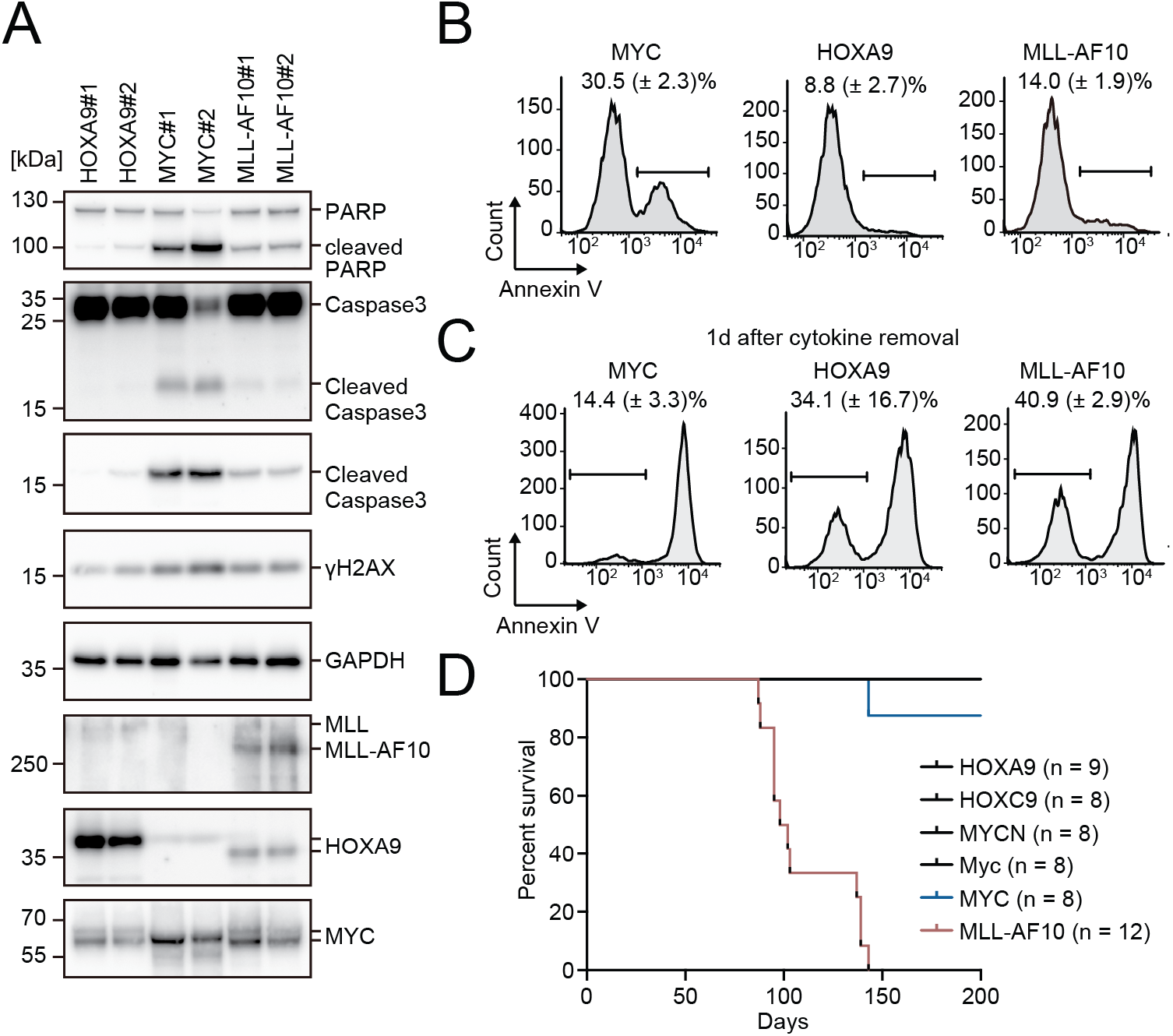
Apoptosis is induced by MYC, while is alleviated by HOXA9 and MLL-AF10. **A**, Protein expression of transgenes and apoptotic markers in HOXA9-, MYC-, and MLL-AF10-ICs. **B** and **C**, Apoptotic tendencies of HOXA9-, MYC-, and MLL-AF10-ICs. Representative FACS plots and the summarized data (Mean with SD, n = 3, biological replicates) of Annexin V staining of HOXA9-, MYC-, and MLL-AF10-ICs in the presence of cytokines (**B**) and 1d after their removal (**C**) are shown. **D**, In vivo leukemogenic potential of MLL target genes and MLL-AF10. Kaplan-Meier curves of mice transplanted with HPCs transduced with the indicated genes and the number of replicates are shown.

Next, we evaluated the leukemogenic potential of HOXA9 and MYC in vivo. Transplantation of HPCs transduced with *MLL-AF10* into syngeneic mice induced leukemia with full penetrance (Figure 3D). In contrast, neither *MYC* nor its homologue *MYCN* was capable of initiating leukemia within 200 days under these experimental conditions, indicating that activation of the MYC high signature alone is insufficient for leukemogenesis in vivo. Although one recipient mouse of *MYC*-transduced HPCs became sick and was sacrificed about 150 days after transplantation, t did not exhibit any leukemia-associated signs. *HOXA9* did not induce leukemia within 200 days either, suggesting that the HOXA9 high signature alone is also insufficient to induce leukemia. Taken together, these results suggest that both high MYC activity and HOXA9-mediated resistance to apoptosis are necessary for driving leukemogenesis in vivo.

### HOXA9 promotes MYC-mediated leukemogenesis

We next examined the gene expression of patients with leukemia using publicly available microarray data of the Microarray Innovations in LEukemia (MILE) study (29). Of 108 cases with MLL translocation, 38 were acute myelogenous leukemia [AML] and 70 were acute lymphoblastic leukemia [ALL]. Most of the cases (78.7%) were categorized in the HOXA9^high^/MEIS1^high^ group, while some were in the HOXA9^low^/MEIS1^high^ (13.0%) or HOXA9^high^/MEIS1^low^ (8.3%) group (Figure 4A). *MYC* was expressed at high levels irrespective of *HOXA9* or *MEIS1* expression. These data indicate variability in transcriptional profiles among *MLL*-rearranged leukemia cases, with *MYC* expression remaining consistently high. HOXA9^low^/MEIS1^high^ leukemia was predominatly found in ALL, likely due to the MLL-AF4 cases, some of which do not express *HOXA* genes (30). HOXA9^high^/MEIS1^low^ leukemia was mainly found in AML (Figure 4B).

**Figure 4.**
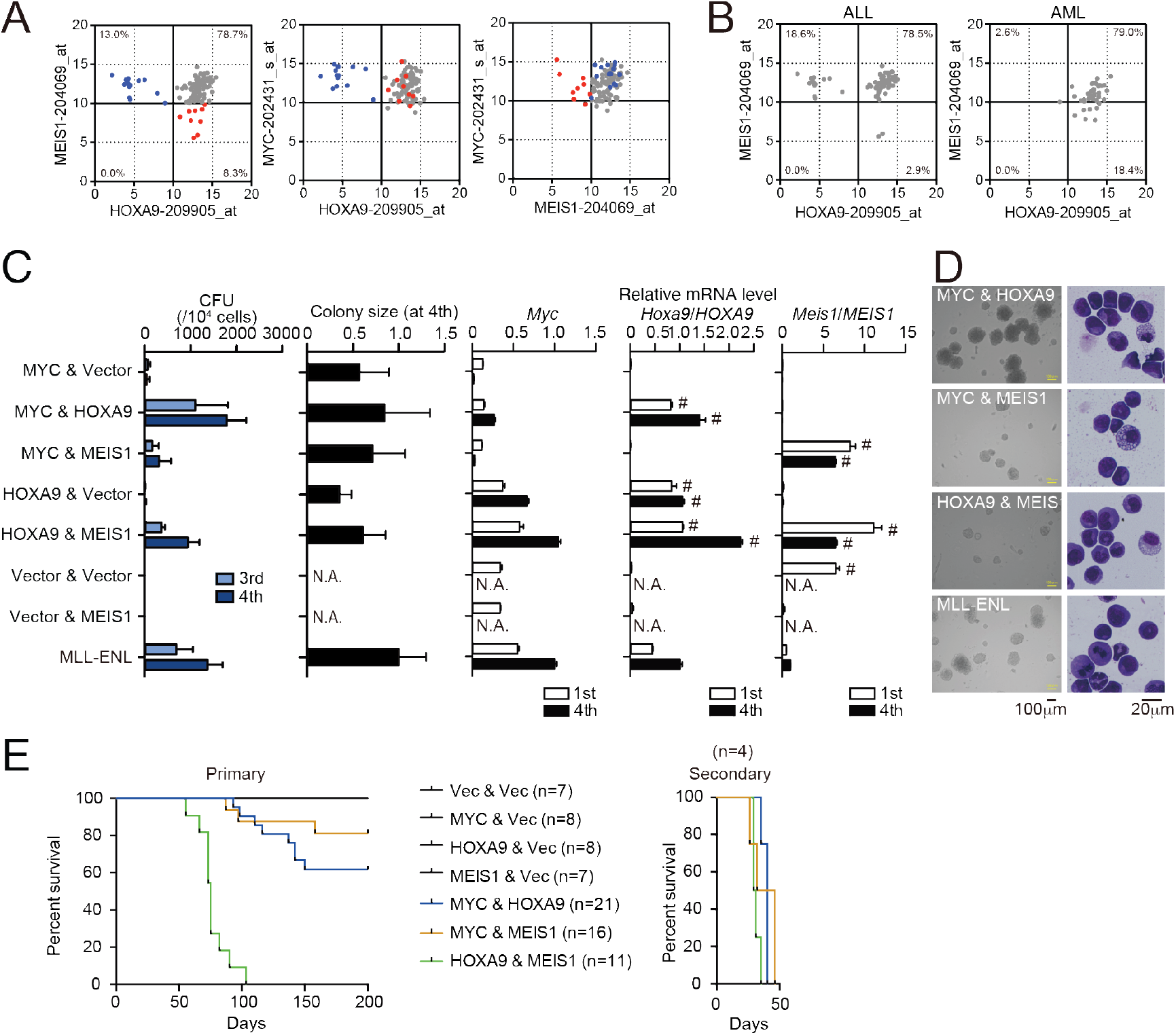
HOXA9 promotes MYC-mediated leukemogenesis. **A** and **B**, Expression profiles of *MLL*-rearranged leukemia patients reported in the MILE study (29). Probe intensities of the indicated genes are plotted for all *MLL*-rearranged leukemia patients (**A**). Patients in the HOXA9^low^/MEIS1^high^ (blue) and HOXA9^high^/MEIS1^low^ (red) groups are highlighted. Probe intensities of *HOXA9* and *MEIS1* are plotted separately by leukemia phenotype (ALL or AML) (**B**). **C**, Transforming potential of various combinations of MLL target genes. CFU (Mean with SD, n = 3, biological replicates) and relative colony size (Mean with SD, n ≥ 100) are shown as in Figure 1C. **D**, Morphologies of the colonies and transformed cells. Bright field (left) and May-Grunwald-Giemsa staining (right) images are shown with scale bars. **E**, In vivo leukemogenic potential of various oncogene combinations. Kaplan-Meier curves of mice transplanted with HPCs transduced with the indicated genes are shown as in Figure 3D. Bone marrow cells from moribund mice were harvested and used for secondary transplantation.

Next, we assessed the effects of combined expression of MYC, HOXA9, and MEIS1 in mouse leukemia models. The MYC/HOXA9 combination exhibited high clonogenicity ex vivo, with colony-forming capacity and colony size comparable to those of MLL-ENL- and HOXA9/MEIS1-ICs (Figure 4C, D). A weak synergy between MYC and MEIS1 was also observed. HOXA9/MEIS1-ICs showed high Myc expression, indicating essential role of Myc in leukemic transformation of HPCs even by this gene set. In the in vivo leukemogenesis assays, MYC, HOXA9, or MEIS1 expression alone did not initiate leukemia, whereas co-expression of HOXA9 and MEIS1 did in all recipient mice as previously reported (Figure 4E) (15). The combined expression of MYC/HOXA9 or MYC/MEIS1 induced leukemia in 38% and 18% of recipient mice, respectively, indicating a synergy of MYC with HOXA9 and MEIS1. qRT-PCR analysis showed that LCs maintained the expression patterns of endogenous *Hoxa9*, *Meis1*, and *Myc* similar to those of the respective ICs (Supplemental Figure 3). We observed rapid onset of leukemia with full penetrance in the secondary transplantation for the three combinations tested, confirming the presence of leukemia-initiating cells (Figure 4E). These results indicate a cooperative role for HOXA9 and MEIS1 in MYC-mediated leukemogenesis with stronger synergy between HOXA9 and MYC.

### HOXA9 functions as a transcription maintenance factor

Some HOXA9 high signature genes, namely *Bcl2*, *Sox4*, and *Igf1*, have been implicated in leukemogenesis (31–34), suggesting that they may be responsible for the synergy between HOXA9 and MYC. Indeed, *Bcl2, Sox4*, and *Igf1* were highly transcribed in HPCs transformed by the combinations containing HOXA9 (i.e. MYC/HOXA9-ICs, HOXA9/vector-ICs, and HOXA9/MEIS1-ICs) but not those lacking it (i.e. MYC/vector-ICs and MYC/MEIS1-ICs)(Figure 5A and Supplemental Figure 4A). Conditional loss of function experiments using HOXA9 conjugated with estrogen receptor (HOXA9-ER) confirmed direct regulation of these genes by HOXA9 (Figure 5B). However, stepwise transduction of *MYC* followed by *HOXA9* failed to upregulate these HOXA9 target genes, indicating that HOXA9 cannot re-activate its target genes once silenced (Figure 5C and Supplemental Figure 4B). These results indicate that HOXA9 is a transcriptional maintenance factor which may be involved in the maintenance of chromatin structure previously activated by other transcriptional/epigenetic factors.

**Figure 5.**
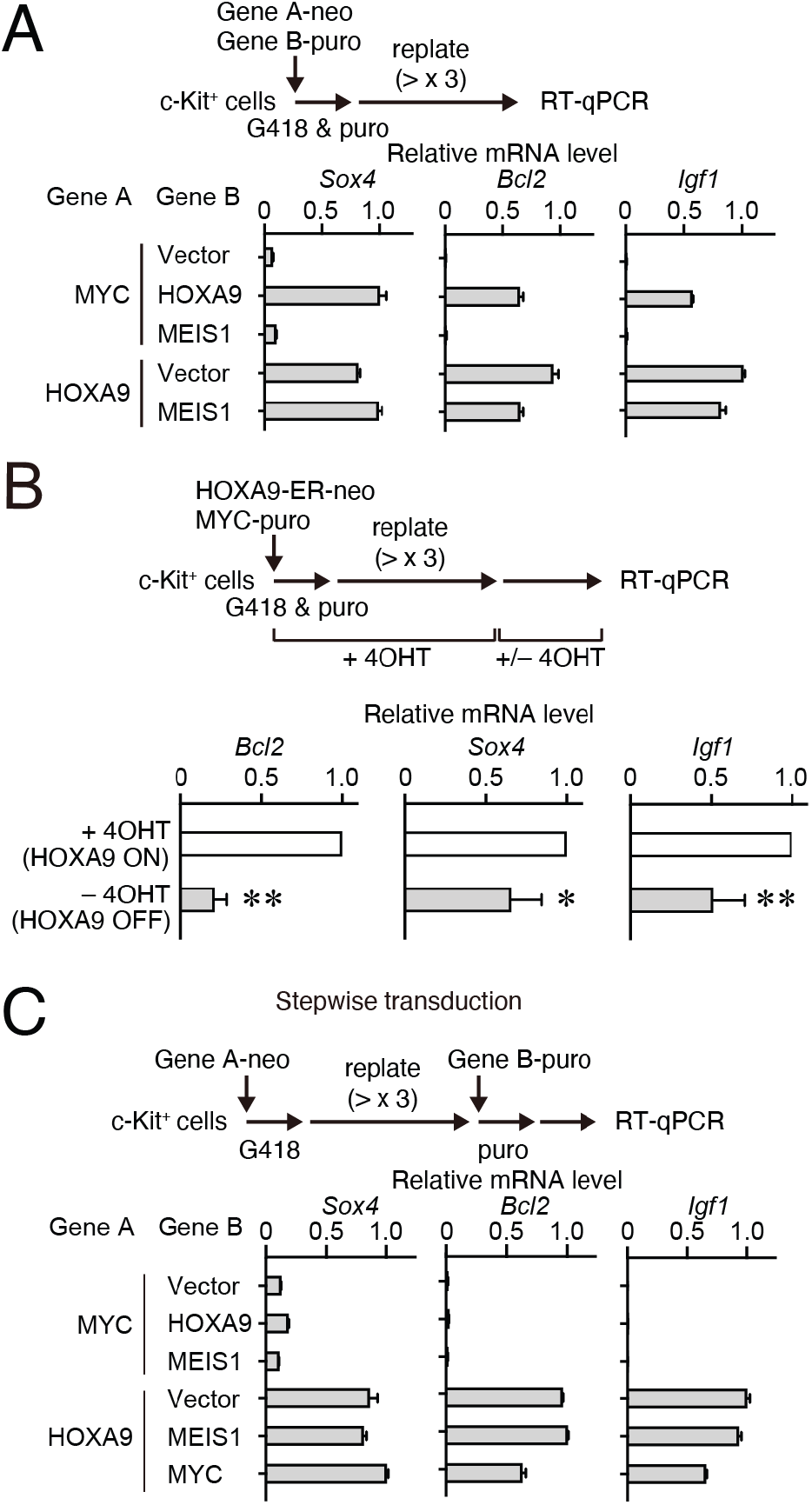
HOXA9 functions as a transcription maintenance factor. **A**. Gene expression of HPCs immortalized by various transgenes. Relative mRNA levels of HOXA9 target genes (Mean with SD, n = 3, PCR replicates) in myeloid progenitors transformed by various combinations of MLL target genes are shown. Two genes were transduced into HPCs in a simultaneous manner. **B**, Gene expression after inactivation of HOXA9. HOXA9-ER and MYC were doubly transduced into HPCs and cultured in the presence of 4-OHT ex vivo. After 4-OHT withdrawal, RT-qPCR analysis was performed for the indicated genes (Mean, n=4, biological replicates). Statistical analysis was performed using unpaired two-tailed Student’s *t*-test. \*\**P* < 0.01, \**P* < 0.05. **C**. Gene expression of HPCs immortalized by step-wise transduction of various transgenes. Relative mRNA levels of HOXA9 target genes in myeloid progenitors transformed by various combinations of MLL target genes are shown as in A. Two genes were transduced into HPCs in a stepwise manner.

### BCL2 and SOX4 promote MYC-mediated leukemogenesis by alleviating apoptosis

To identify the roles for BCL2 and SOX4 in leukemic transformation, we evaluated the leukemogenic potential of combined expression of BCL2 or SOX4 with MYC. In myeloid progenitor transformation assays, SOX4 by itself showed weak immortalization capacity as previously reported (31), while BCL2 did not transform HPCs (Figure 6A). Co-expression of BCL2 or SOX4 with MYC led to a substantial increase in colony-forming capacity. The combined expression of BCL2 with MYC induced leukemia in vivo with a penetrance similar to that with the MYC/HOXA9 combination (Figures 4E and 6B) in accord with previous reports (35, 36). SOX4 also promoted MYC-mediated leukemogenesis in vivo (Figure 6B), while neither MYC, BCL2, nor SOX4 alone induced leukemia within 200 days. It should be noted that single gene transduction of MYC and SOX4 was shown to induce leukemia by others in different settings albeit low penetrance (32, 36). These differences are possibly due to the differences of virus titers and viral genome integration-related gene activation. Furthermore, the combined expression of HOXA9, BCL2, or SOX4 with MYC alleviated apoptotic tendencies (Figure 6C, D). Although SOX4 is reported to modulate transcription of pro/anti-apoptotic genes (37), their expression was not drastically altered by SOX4 in this context (Figure 6E and F). Thus, the mechanism underlying the anti-apoptotic properties of SOX4 in this setting is currently unclear. Taken together, the results indicate that multiple anti-apoptotic pathways mediated by BCL2 and SOX4 promote MYC-mediated leukemic transformation.

**Figure 6.**
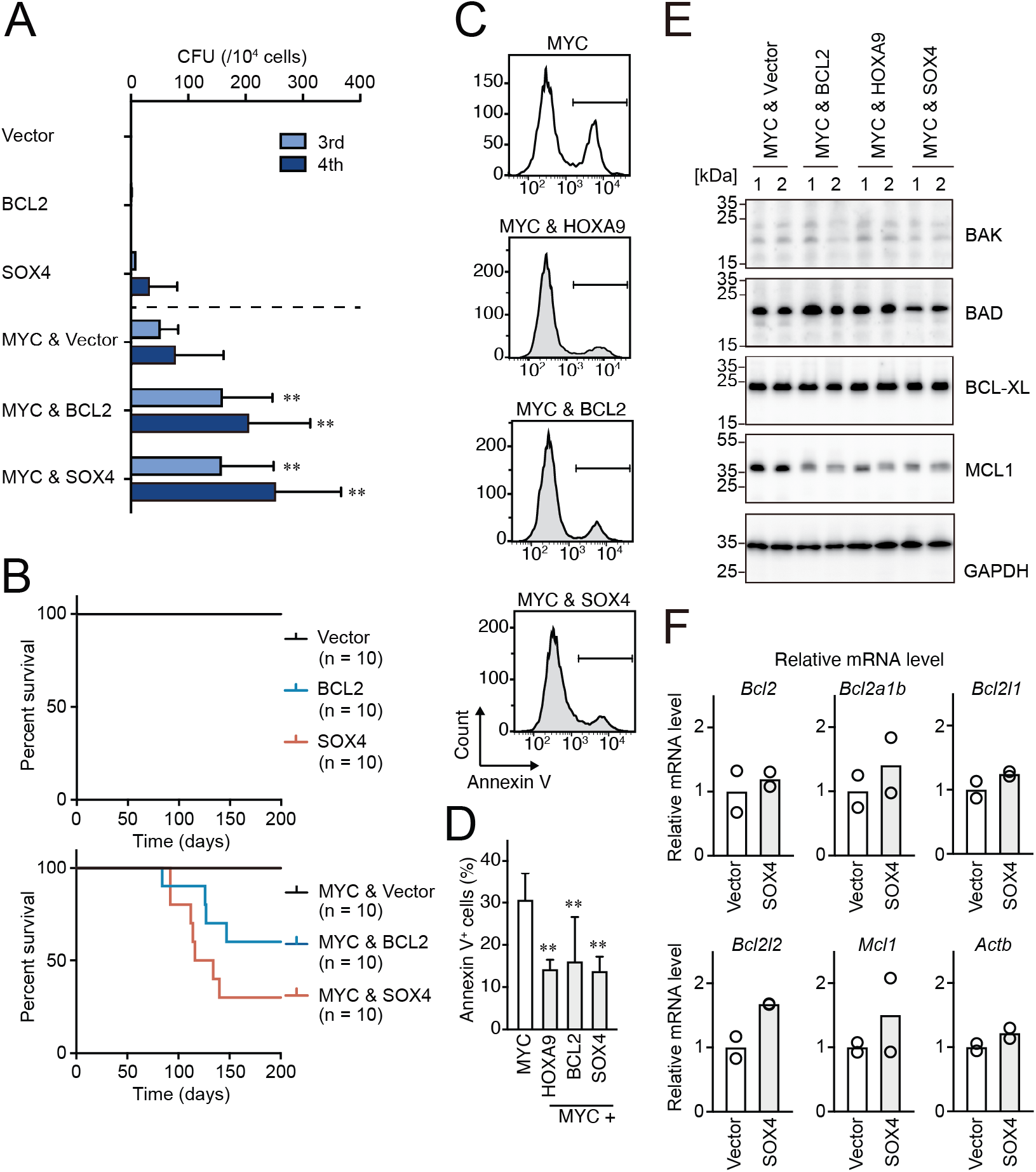
BCL2 and SOX4 promote MYC-mediated leukemogenesis by alleviating apoptosis. **A**, Transforming potential of various combinations of MYC and HOXA9 target genes. CFU (Mean with SD, n = 3, biological replicates) is shown as in Figure 1C. **B**, In vivo leukemogenic potential of various combinations of MYC and HOXA9 target genes. Kaplan-Meier curves of mice transplanted with HPCs transduced with the indicated genes are shown as in Figure 3D. **C** and **D**, Apoptotic tendencies of *MYC*-expressing progenitors co-transduced with *HOXA9*, *BCL2*, or *SOX4*. Representative FACS plots (**C**) and the summarized data (**D**) (Mean with SD, n = 3, biological replicates) of Annexin V staining are shown. Statistical analysis was performed using ordinary one-way ANOVA with MYC-ICs. **E**, Expression of apoptosis-related proteins in HPCs transformed by various combinations of transgenes. Western blots of HPCs transformed by indicated transgenes are shown **F**, Relative expression levels of apoptosis-related genes in MYC/SOX4-ICs and MYC/vector-ICs. Relative mRNA levels of the indicated apoptosis-related genes (Mean, n = 2, biological replicates) are shown.

### Endogenous BCL2 and SOX4 support the initiation and maintenance of leukemia

To examine the roles for endogenous BCL2 and SOX4 in the initiation and maintenance of LCs, we conducted myeloid progenitor transformation and in vivo leukemogenesis assays using *Bcl2*- and *Sox4*-knockout HPCs. We transduced various oncogenes into HPCs isolated from fetal livers of *Bcl2*- and *Sox4*-knockout embryos (38, 39) and cultured ex vivo. Neither *Bcl2* nor *Sox4* deletion affected the proliferation of HPCs transduced with *HOXA9*, *MYC*, or *MLL-AF10*, indicating that BCL2 and SOX4 are dispensable for proliferation ex vivo (Supplemental Figure 5A). On the other hand, *Bcl2* deletion delayed the onset of leukemia induced by MLL-AF10 and the HOXA9/MEIS1 combination in vivo (Figure 7A). *Sox4* deletion also delayed the onset of HOXA9/MEIS1-induced leukemia, although it did not affect MLL-AF10-induced leukemia (Figure 7B). These results indicate that endogenous BCL2 and SOX4 partially contribute to initiate leukemia in vivo.

**Figure 7.**
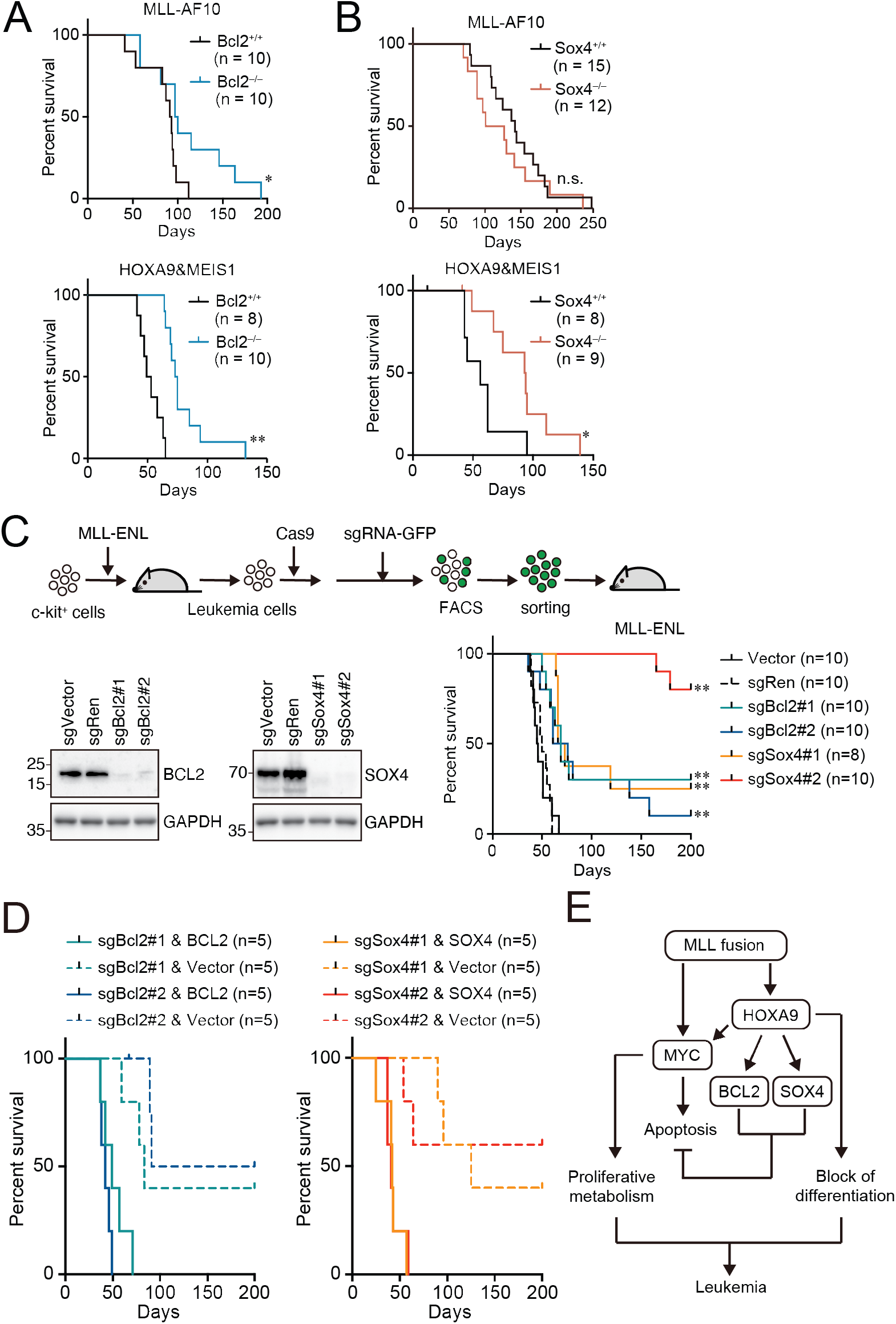
Endogenous BCL2 and SOX4 support the initiation and maintenance of leukemia. **A** and **B**, Effects of *Bcl2*- or *Sox4*-deficiency on the initiation of leukemogenesis in vivo. HPCs were isolated from *Bcl2*^+/+^ or *Bcl2*^−/−^ (**A**) or *Sox4*^+/+^ or *Sox4*^−/−^ (**B**) embryos and transduced with *MLL-AF10* or the *HOXA9*/*MEIS1* combination. Kaplan-Meier curves of mice transplanted with the transduced HPCs are shown. Statistical analysis was performed using the log-rank test and Bonferroni correction with the wildtype control. \**P* ≤ 0.05. **C**, Effects of *Bcl2*- or *Sox4*-deficiency on the maintenance of leukemia initiating cells. Western blots of MLL-ENL leukemia cells transduced with sgRNA for BCL2 or SOX4 are shown on the left. Kaplan-Meier curves of mice transplanted with MLL-ENL leukemia cells transduced with the indicated sgRNAs are shown on the right. Statistical analysis was performed using the log-rank test and Bonferroni correction with the vector control. **D**, Rescue of in vivo leukemogenic potential by sgRNA-resistant transgenes. Before transduction of sgRNA, MLL-ENL-LCs were transduced with sgRNA-resistant *BCL2* or *SOX4*. In vivo leukemogenesis assay was performed as described in Figure 3D. **E**, A model illustrating HOXA9-mediated pathogenesis in *MLL*-rearranged leukemia.

Next, we examined the roles for endogenous BCL2 and SOX4 in the maintenance of leukemia initiating cells using a mouse leukemia cell line that we previously established with MLL-ENL (23). CRISPR/Cas9-mediated sgRNA competition assays indicated negligible contribution of *Bcl2* and *Sox4* to MLL-ENL-LC proliferation ex vivo (Supplemental Figure 5B). To test their roles in vivo, we transplanted *Bcl2*- or *Sox4*-deficient LCs into syngeneic mice, where we observed delayed onset and reduced incidence rate for both *Bcl2*- and *Sox4*-knockout LCs (Figure 7C). Forced expression of sgRNA-resistant cDNAs of *BCL2* and *SOX4* restored leukemogenic potential, validating the on-target effects of sgRNAs (Figure 7D). These results suggest that MLL-ENL-LCs are partially dependent on endogenous BCL2 and SOX4 in vivo. Taken together, our results indicate that *MLL*-rearranged leukemia cells depend on the expression of multiple anti-apoptotic genes via HOXA9 to varying degrees to achieve survival advantages for disease initiation and maintenance (Figure 7E).

## Discussion

In this study, we found that HOXA9 regulates a variety of genes to maintain hematopoietic precursor identity and its associated anti-apoptotic properties. In leukemic transformation, MLL fusion proteins exploit both HOXA9 and MYC downstream pathways. Accordingly, HOXA9 and MYC synergistically induce leukemia in mouse models. Thus, we propose that MLL fusion proteins employ two arms to promote oncogenesis: MYC-mediated proliferation and HOXA9-mediated resistance to differentiation/apoptosis.

It is widely accepted that two types of mutations need to occur before leukemia onset; class I mutations that confer proliferative advantages and class II mutations that block differentiation (40). However, it was unclear how MLL mutations fit this theory because *MLL*-rearranged leukemia does not require additional mutations in many cases (41). A comparison of gene expression profiles of HOXA9- and MYC-transformed HPCs demonstrated that HOXA9 maintains the expression of a wide range of genes associated with hematopoietic precursor identity. Thus, HOXA9 appears to maintain intrinsic transcriptional programs of immature HPCs, which are programed to be silenced during differentiation. On the other hand, MYC upregulates genes involved with de novo protein synthesis or metabolism. This is in line with the known involvement of MYC in proliferation and the associated anabolic processes (42). Furthermore, MLL-AF10 activated both the HOXA9 high and MYC high signature genes to induce leukemia while ectopic expression of HOXA9 and MYC synergistically induced leukemia. Thus, our findings suggest that MLL fusion proteins activate both HOXA9- and MYC-dependent programs as alternative mechanisms to the combination of class I and II mutations.

Therapeutic efficacy of BCL2 inhibitor has been reported in AML including MLL leukemia (43). Because there is a correlation between the expression levels of HOX proteins and the sensitivity to BCL2 inhibitor in AML patient samples, the aberrant expression of HOXA9 is the potential mechanisms for BCL2-dependence of MLL leukemia (44, 45). Oncogenic MYC expression often leads to apoptosis, which need to be alleviated by additional genetic events for leukemic cell survival (28). Indeed, co-expression of HOXA9, BCL2 or SOX4 promoted MYC-mediated leukemogenesis, while MYC alone was insufficient to induce leukemia in vivo. Although the downstream mechanisms could not be addressed, SOX4 also exhibited anti-apoptotic effects on MYC-expressing cells. Importantly, there was a partial effect of single gene knockout of *Bcl2* and *Sox4* on leukemia initiation and maintenance. This indicates that multiple anti-apoptotic pathways are exploited by MLL fusion proteins and that blocking a single anti-apoptotic pathway may be insufficient to completely abrogate leukemic potential. Thus, simultaneously blocking multiple anti-apoptotic pathways may be required for efficient molecularly targeted therapy of *HOXA9*-expressing leukemia.

Our results also provide an insight into the mode of function of HOXA9. *HOX* genes are known to express in a position-specific manner, conferring a positional identity to a cell. For example, similarly functional fibroblasts derived from different parts of a body express different *HOX* genes (5). Thus, it is unlikely that HOXA9 functions as a major upstream factor which determines tissue-specific gene expression by turning a silenced chromatin into transcriptionally active chromatin. HOX proteins likely play a supportive role to maintain gene expression which was activated by other transcriptional regulators. Our observation that HOXA9 cannot reactivate gene expression once silenced fits to this hypothesis. Accordingly, HOXA9 maintains a subset of genes related to hematopoietic identity when expressed in hematopoietic precursors. Recently, it has been reported that HOXA9 may recruit enhancer apparatuses, as it colocalizes with active enhancer mark (i.e. acetylated histone H3 lysine 27). We speculate that HOXA9 may support the maintenance of an active enhancer, but unlikely establishes it on a silenced chromatin. Further functional analysis of HOXA9 is required to understand how HOXA9 regulates gene expression.

In summary, our results describe the oncogenic roles for HOXA9 as transcriptional maintenance factor for multiple anti-apoptotic genes, which are necessary to promote MYC-mediated leukemogenesis. In case of *MLL*-rearranged leukemia, MLL fusion proteins directly activate both *MYC* and *HOXA9*, while HOXA9 maintains expression of *MYC*, *BCL2*, and *SOX4*, achieving high MYC activity and anti-apoptotic properties simultaneously (Figure 7E). Thus, *MLL*-rearranged leukemia cells acquired highly proliferative potentials and survival advantages at the same time, using HOXA9 as a key mediator.

## Materials and Methods

### Vector constructs

For protein expression vectors, cDNAs obtained from Kazusa Genome Technologies Inc. (46) were modified by PCR-mediated mutagenesis and cloned into the pMSCV vector (for virus production) or pCMV5 vector (for transient expression) by restriction enzyme digestion and DNA ligation (Supplemental Table 1). The MSCV-neo MLL-ENL, and MLL-AF10 vectors have been previously described (23). sgRNA-expression vectors were constructed using the pLKO5.sgRNA.EFS.GFP vector (47). shRNA-expression vectors were purchased from Dharmacon. The target sequences are listed in Supplemental Table 2 (sgRNA) and Supplemental Table 3 (shRNA).

### Cell lines

HEK293T cells were a gift from Michael Cleary and were authenticated by the JCRB Cell Bank in 2019 (Supplemental Table 1). HEK293TN cells were purchased from System Biosciences. The cells were cultured in Dulbecco’s modified Eagle’s medium (DMEM) supplemented with 10% fetal bovine serum (FBS) and penicillin-streptomycin (PS). The Platinum-E (PLAT-E) ecotropic virus packaging cell line—a gift from Toshio Kitamura (48)—was cultured in DMEM supplemented with 10% FBS, puromycin, blasticidin, and PS. The human leukemia cell line HB1119, also a gift from Michael Cleary (49), was cultured in RPMI 1640 medium supplemented with 10% FBS and PS. MLL-ENL leukemia cells (MLL-ENLbm0713) were described previously (23). Cells were incubated at 37 °C and 5% CO_2_.

### Animal models

For the in vivo leukemogenesis assay, 8-week-old female C57BL/6JJcl (C57BL/6J) or C. B-17/Icr-*scid/scid*Jcl (SCID) mice were purchased from CLEA Japan (Tokyo, Japan). *Bcl2*-knockout mice were a gift from Yoshihide Tsujimoto and provided via RIKEN BRC (38). *Sox4*-knockout mice were a gift from Hans Clevers and provided via RIKEN BRC (39).

### Western blotting

Western blotting was performed as previously described (50). Antibodies used in this study are listed in Supplemental Table 1.

### Virus production

Ecotropic retrovirus was produced using PLAT-E packaging cells (48). Lentiviruses were produced in HEK293TN cells using the pMDLg/pRRE, pRSV-rev, and pMD2.G vectors, all of which were gifts from Didier Trono (51). The virus-containing medium was ha0,rvested 24–48 h following transfection and used for viral transduction.

### Myeloid progenitor transformation assay

The myeloid progenitor transformation assay was carried out as previously described (26, 27). Bone marrow cells were harvested from the femurs and tibiae of 5-week-old female C57BL/6J mice. c-Kit^+^ cells were enriched using magnetic beads conjugated with an anti-c-Kit antibody (Miltenyi Biotec), transduced with a recombinant retrovirus by spinoculation, and then plated (4 × 10^4^cells/ sample) in a methylcellulose medium (Iscove’s modified Dulbecco’s medium, 20% FBS, 1.6% methylcellulose, and 100 μM β-mercaptoethanol) containing murine stem cell factor (mSCF), interleukin 3 (mIL-3), and granulocyte-macrophage colony-stimulating factor (mGM-CSF; 10 ng/mL each). During the first culture passage, G418 (1mg/mL) or puromycin (1μg/mL) was added to the culture medium to select for transduced cells. *Hoxa9* expression was quantified by qRT-PCR after the first passage. Cells were then re-plated once every 4-6 days with fresh medium; the number of plated cells for the second, third, and fourth passages was 4 × 10^4^, 2 × 10^4^, and 1 × 10^4^ cells/well, respectively. CFUs were quantified per 10^4^ plated cells at each passage.

### In vivo leukemogenesis assay

In vivo leukemogenesis assays were carried out as previously described (26, 52). c-Kit^+^ cells (2 × 10^5^) prepared from the femurs and tibiae of 5-week-old female C57BL/6J mouse were transduced with retrovirus by spinoculation and intravenously transplanted into sublethally irradiated (5–6 Gy) C57BL/6J mice. For secondary leukemia, leukemia cells (2 × 10^5^) cultured ex vivo for more than three passages were transplanted. As for knockouts of *Bcl2* and *Sox4*, mice heterozygous for *Bcl2* or *Sox4* were crossed, and c-Kit^+^ cells were isolated from fetal livers at E14–15 (for Bcl2) or E13 (for Sox4). The next day, cells were transduced with *MLL-AF10* or *HOXA9*/*MEIS1* and transplanted intravenously into sublethally irradiated (2.5 Gy) SCID mice [2 × 10^5^ (for *Bcl2*) or 1 × 10^5^ cells/mouse (for *Sox4*)].

### qRT-PCR

Total RNA was isolated using the RNeasy Mini Kit (Qiagen) and reverse-transcribed using the Superscript III First Strand cDNA Synthesis System (Thermo Fisher Scientific) with oligo (dT) primers. Gene expression was analyzed by qPCR using TaqMan probes (Thermo Fisher Scientific). Relative expression levels were normalized to those of *GAPDH*/*Gapdh* or *TBP*/*Tbp* and determined using a standard curve and the relative quantification method, according to manufacturer’s instructions (Thermo Fisher Scientific). Commercially available PCR probes used are listed in Supplemental Table 1.

### ChIP-qPCR and ChIP-seq

The eluted material obtained by fanChIP was extracted by phenol/chloroform/isoamyl alcohol. DNA was precipitated with glycogen, dissolved in TE buffer, and analyzed by qPCR (ChIP-qPCR) or deep sequencing (ChIP-seq). The qPCR probe/primer sequences are listed in Supplemental Table 4. Deep sequencing was performed using the TruSeq ChIP Sample Prep Kit (Illumina) and HiSeq2500 (Illumina) at the core facility of Hiroshima University and described in our previous publication (23).

### RNA-seq

Total RNA was prepared using the RNeasy Kit (Qiagen) and analyzed using a Bioanalyzer (Agilent Technologies). Deep sequencing was performed using a SureSelect Strand Specific RNA Library Prep Kit (Agilent Technologies) and HiSeq2500 (Illumina) with 51-bp single-end reads at the core facility of Hiroshima University. Sequenced reads were mapped to the mouse genome assembly mm9 using TopHat 2.0.14 (53) and read counts were normalized with Cufflinks 2.2.1 2010 (54). Data were trimmed by removing lowly expressed genes whose FKPM values were less than 2, after which the top 50 HOXA9 high and MYC high signature genes were visualized as a heatmap using the Complex Heatmap package (55). GSEA analysis and KEGG pathway analysis was performed on the top 500 genes of the HOXA9 or MYC high signature using the DAVID website (56).

### sgRNA competition assay

*Cas9* was introduced via lentiviral transduction using the pKLV2-EF1a-Cas9Bsd-W vector (57). Cas9-expressing stable lines were established with blasticidin (10–30 μg/mL) selection. The targeting sgRNA was co-expressed with GFP via lentiviral transduction using pLKO5.sgRNA.EFS.GFP vector (47). Percentages of GFP^+^ cells were initially determined by FACS analysis at 2 or 3 days after sgRNA transduction, and then measured once every 3–5 days.

### FACS analysis and sorting

To detect apoptosis, two to five million cells were suspended in 200 μL of reaction buffer (140 mM NaCl, 10 mM HEPES, 2.5 mM CaCl_2_, and 0.1% BSA) and incubated with APC-Annexin V for 15 min and PE-PI for 5 min at room temperature. The cells were then centrifuged, resuspended in fresh reaction buffer, and analyzed with FACS Melody (BD Bioscience). FACS sorting of mouse bone marrow cells was performed with fluorophore-conjugated antibodies listed in Supplemental Table 1 as previously described (58).

### Accession numbers

Deep sequencing data used in this study have been deposited in the DNA Data Bank of Japan (DDBJ) Sequence Read Archive under the accession numbers listed in Supplemental Table 5 (ChIP-seq) and Supplemental Table 6 (RNA-seq).

### Statistics

Statistical analysis was performed using GraphPad Prism 7 software. Data are presented as the mean with standard deviation (SD). Comparisons between two groups were analyzed by unpaired two-tailed Student’s *t*-test, while multiple comparisons were performed by ordinary one-way ANOVA followed by Dunnett’s test or two-way ANOVA. Mice transplantation experiments were analyzed by the log-rank test and Bonferroni correction was applied for multiple comparisons. *P* values < 0.05 were considered statistically significant. n.s.: *P*>0.05, *: *P* ≤ 0.05, **: *P* ≤ 0.01, ***: *P* ≤ 0.001, and ****: *P* ≤ 0.0001.

### Study approval

All animal experimental protocols were approved by the National Cancer Center (Tokyo Japan) Institutional Animal Care and Use Committee.

## Acknowledgments

We thank Yuzo Sato, Makiko Okuda, Megumi Nakamura, Etsuko Kanai, Aya Nakayama, Boban Stanojevic, and Ayako Yokoyama for technical assistance. We thank Drs. Yoshihide Tsujimoto and Hans Clevers for providing us the knockout mouse lines of *Bcl2* and *Sox4*, respectively. We also thank all members of the Shonai Regional Industry Promotion Center for their administrative support. This work was supported by the Japan Society for the Promotion of Science (JSPS) KAKENHI grants (16H05337 and 19H03694 to A.Y.; 19K16791 to R.M.) and in part by research funds from the Yamagata prefectural government, the City of Tsuruoka, Dainippon Sumitomo Pharma Co. Ltd., and the Friends of Leukemia Research Fund.

## Author contributions

R.M., H.O., S.T., Y.S., and A.Y. performed the experiments. A.K., H.M., and T.I. performed deep sequencing. A.K. analyzed deep sequencing data. A.Y. conceived of the project. A.Y. and R.M. wrote the paper.

## Supplemental information

**Supplemental Figure 1.**
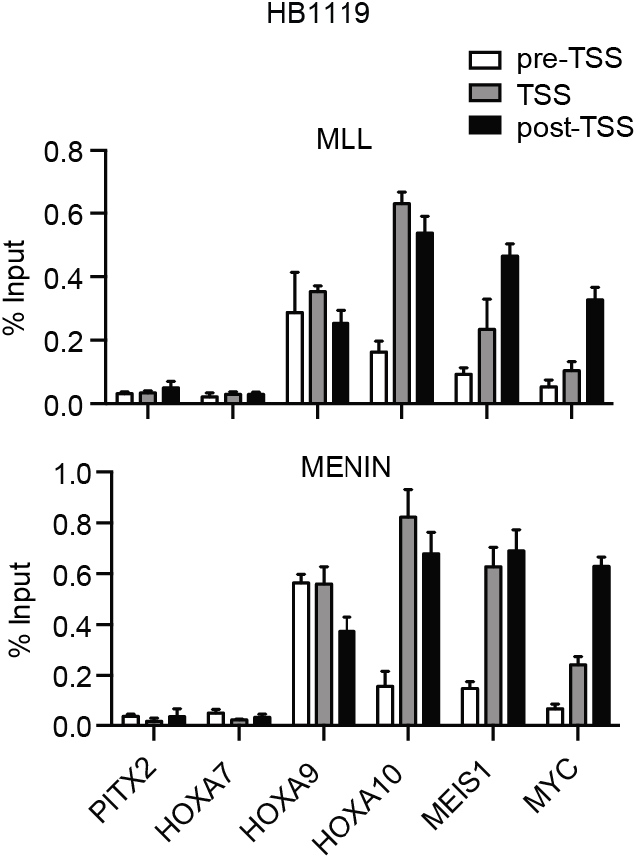
Association of the MLL-ENL complex on target promoters. Genomic localization of the MLL-ENL complex in HB1119 cells. ChIP-qPCR was performed using anti-MLL and MENIN antibodies. Precipitated DNA was subjected to qPCR using specific probes for the pre-TSS (−1.0 to −0.5 kb from the TSS), TSS (0 to 0.5 kb from the TSS), and post-TSS (1.0 to 1.5 kb from the TSS) regions of the indicated genes. ChIP signals were expressed as the percent input (Mean with SD, n = 3, PCR replicates).

**Supplemental figure 2.**
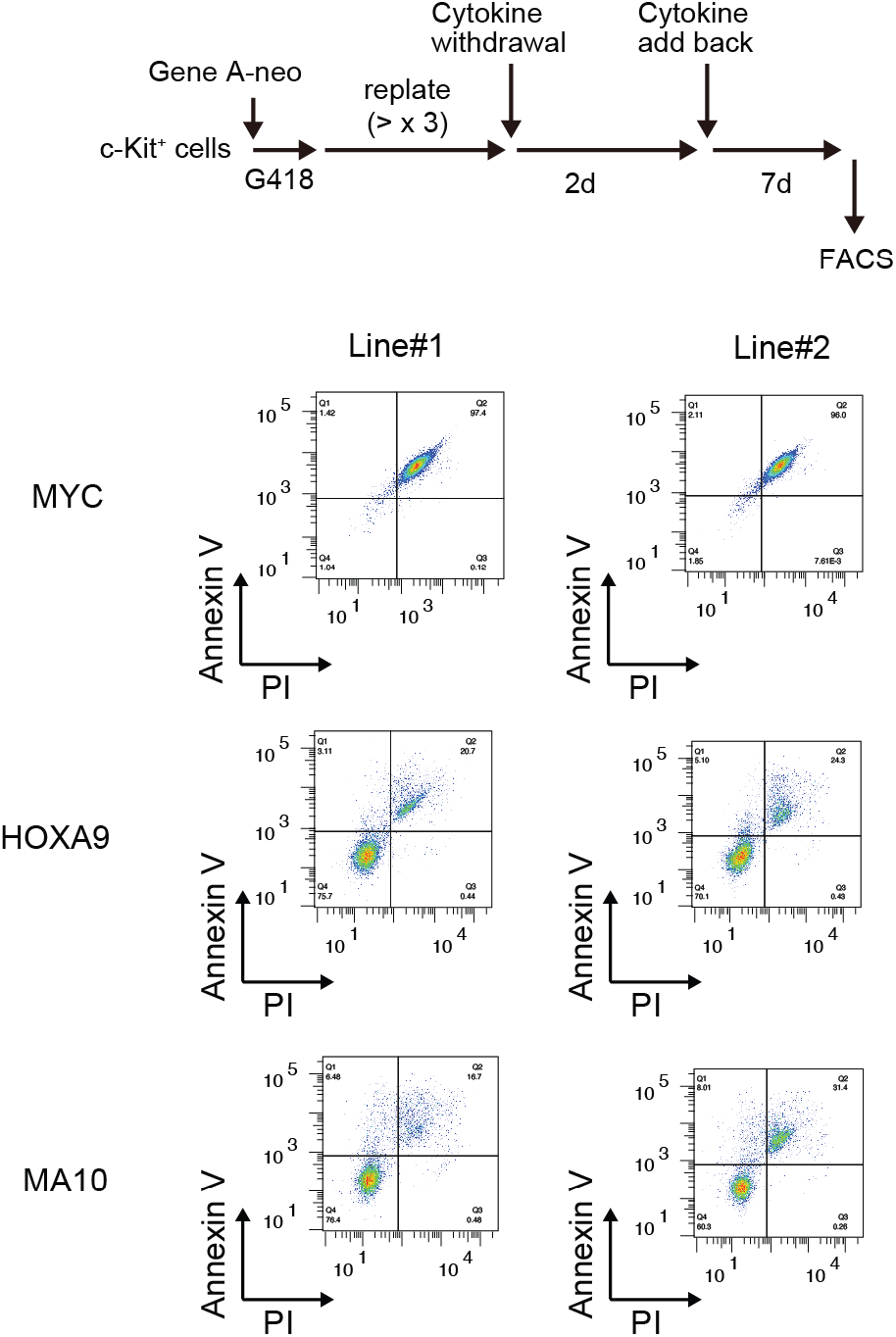
Recovery of HOXA9-expressing cells from cytokine withdrawal. MYC-, HOXA9-, MLL-AF10-ICs were subjected to cytokine withdrawal for 2 d, then cultured in the presence of cytokines for 7d, and examined by FACS for Annexin V and PI.

**Supplemental figure 3.**
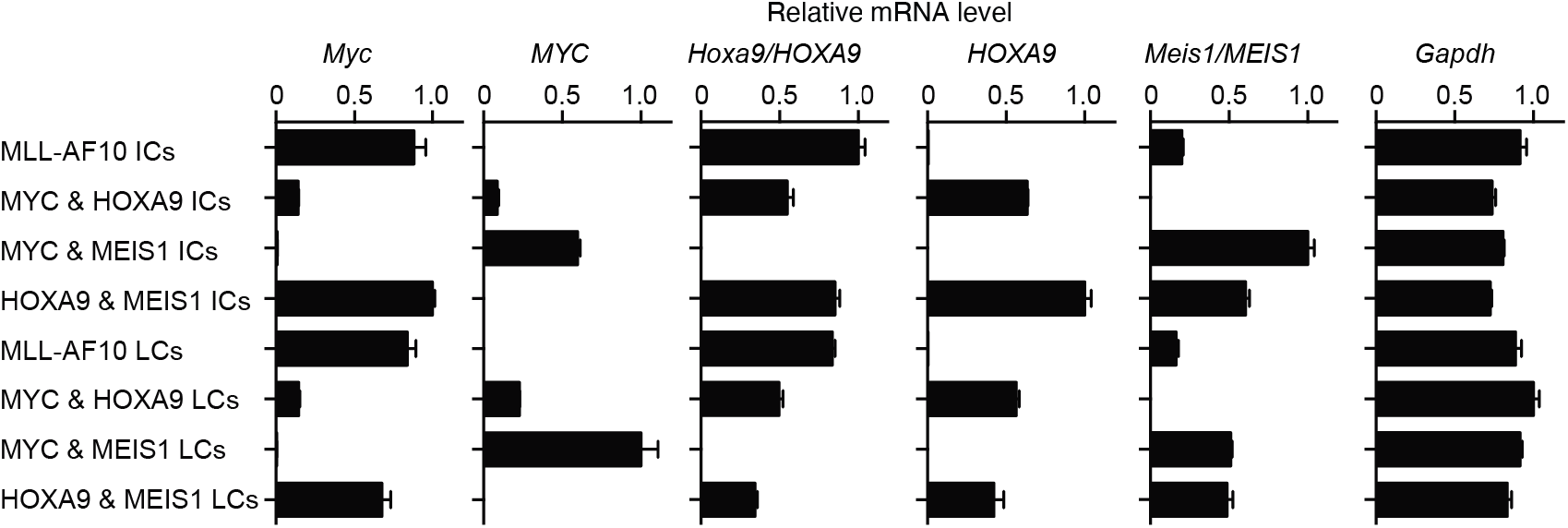
Expression profiles of the LCs established by various gene combinations. LCs were harvested from moribund mice during the transplantation assay as described in Figure 4E. Relative mRNA levels of the indicated genes (Mean with SD, n=3, PCR replicates) are shown along with those of the respective ICs.

**Supplemental figure 4.**
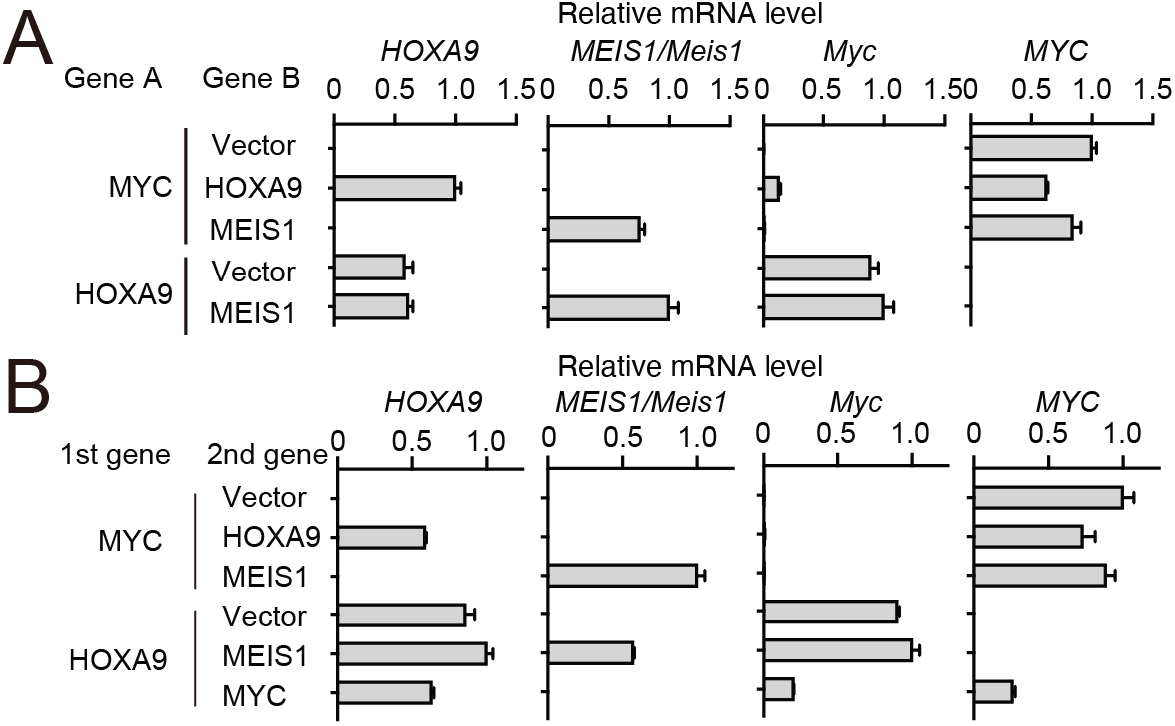
Gene expression of hematopoietic progenitors transformed by various combinations of transgenes. **A** and **B**, Relative mRNA levels of the transgenes and endogenous *Myc* in the cells used in Figure 5A (**A**) and 5C (**B**).

**Supplemental figure 5.**
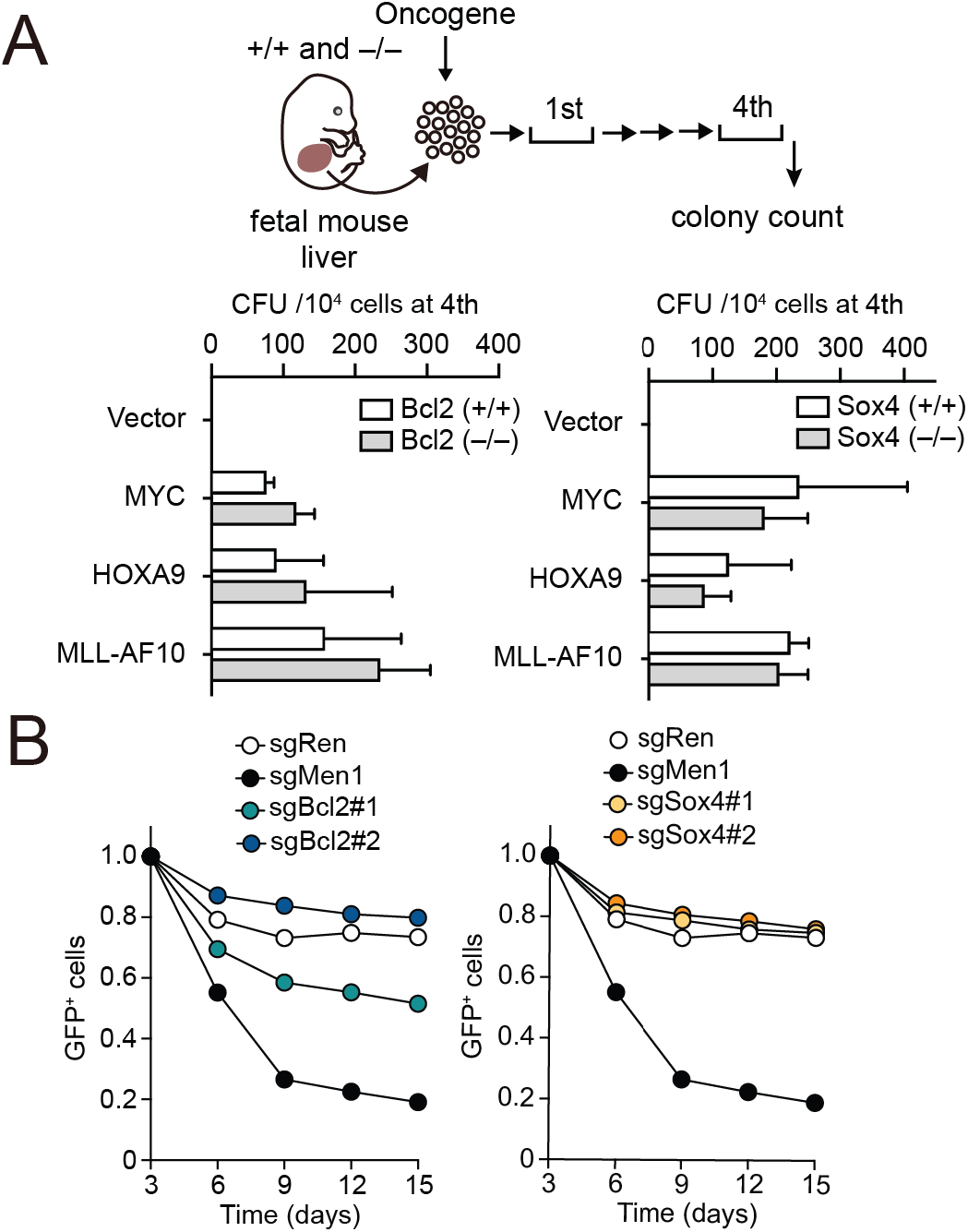
Roles for endogenous BCL2 and SOX4 in transformation ex vivo. **A**, Requirement of BCL2 and SOX4 in transformation ex vivo. Empty vector, *MLL-AF10*, *HOXA9*, or *MYC* were transduced into HPCs isolated from fetal liver of the indicated genotypes. CFU (Mean with SD, n = 3, biological replicates) of myeloid progenitors derived from *Bcl2*- or *Sox4*-knockout embryos is shown. **B**, Effects of *Bcl2* or *Sox4* knockout on MLL-ENL-leukemia cells. sgRNA/GFP was transduced into Cas9-expressing MLL-ENL-LCs. Changes in GFP^+^ cells (%) (Mean n=3, biological replicates) are shown with sg*Men1* serving as a positive control.

**Supplemental Table 1.**
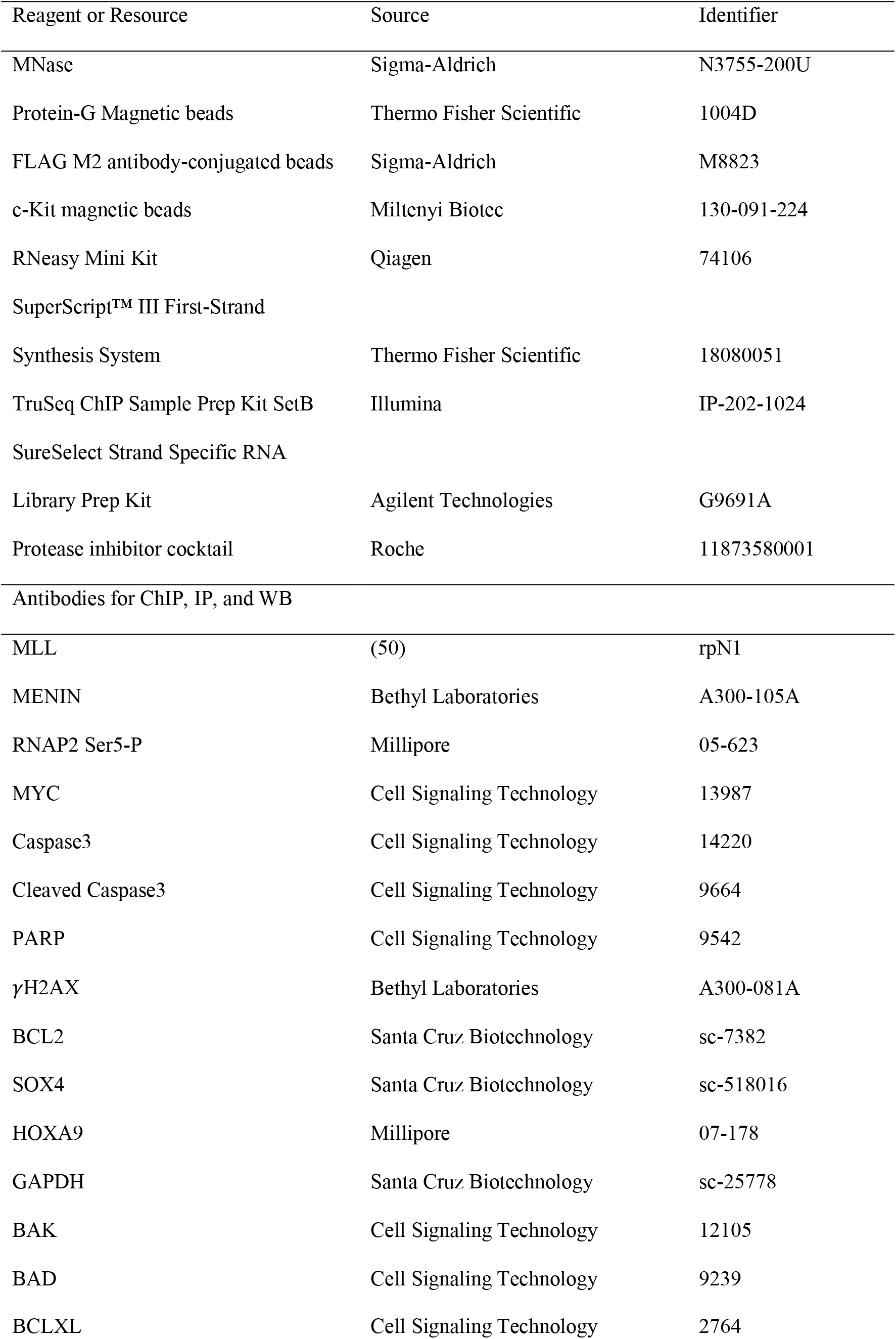
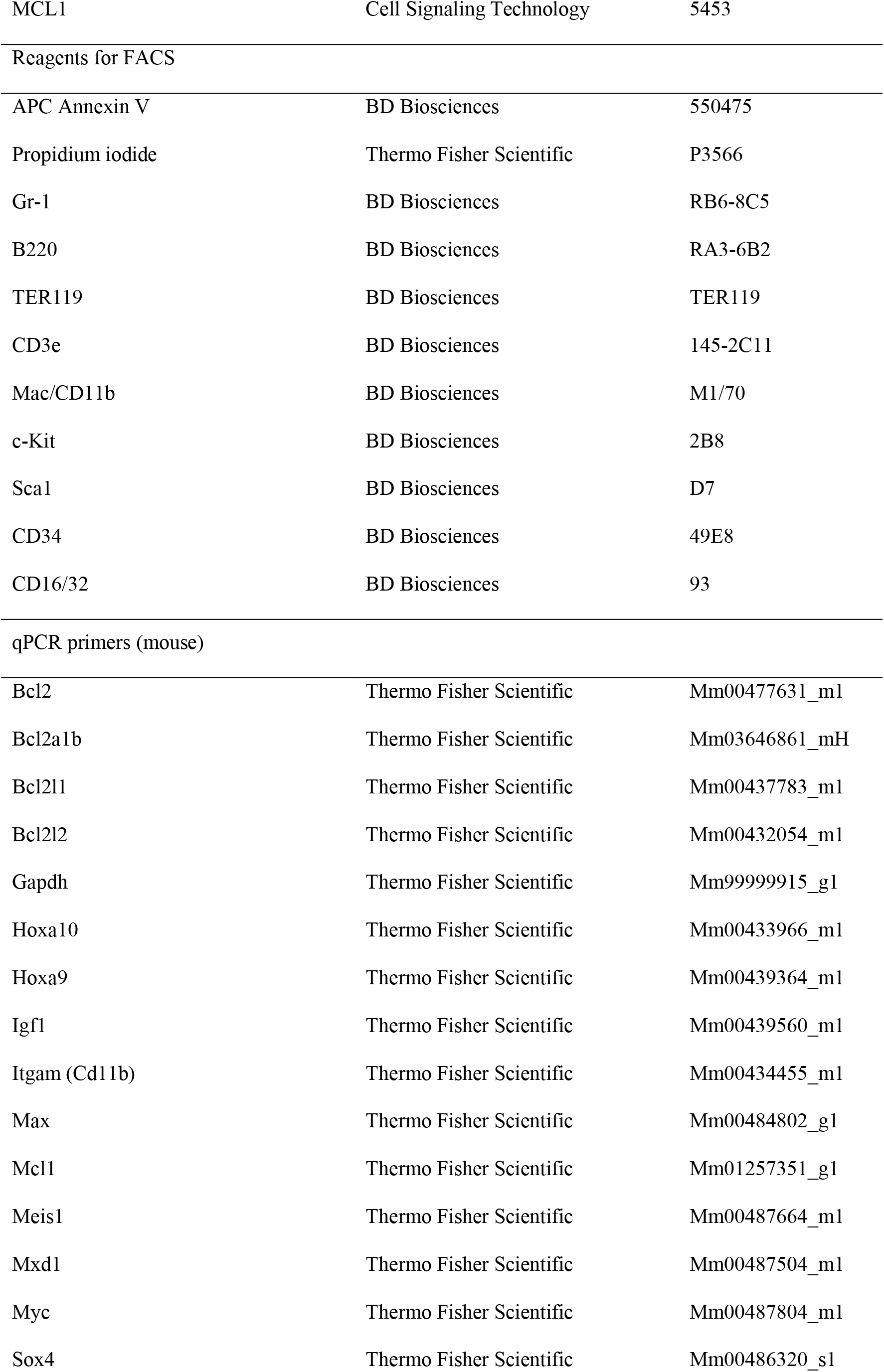
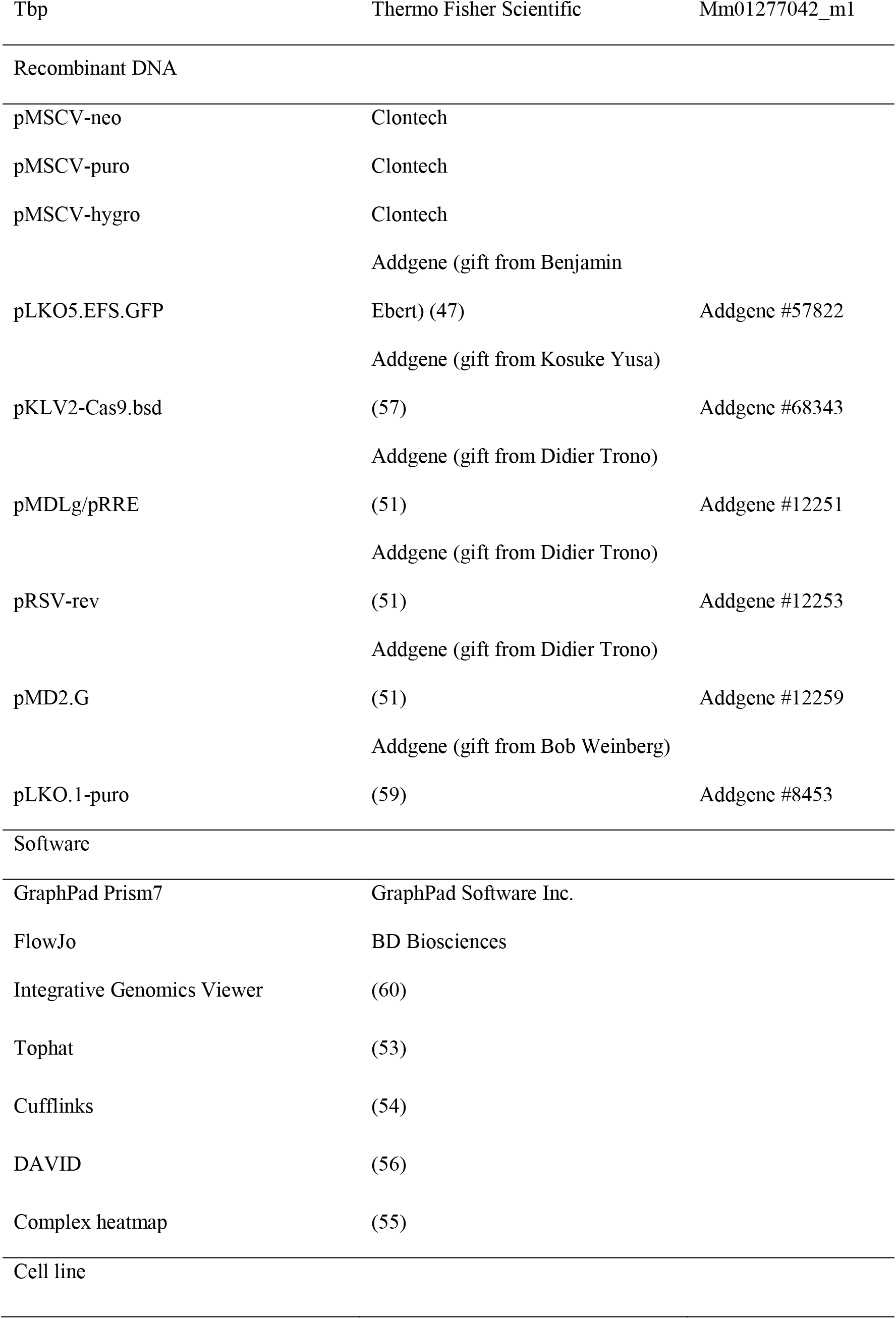
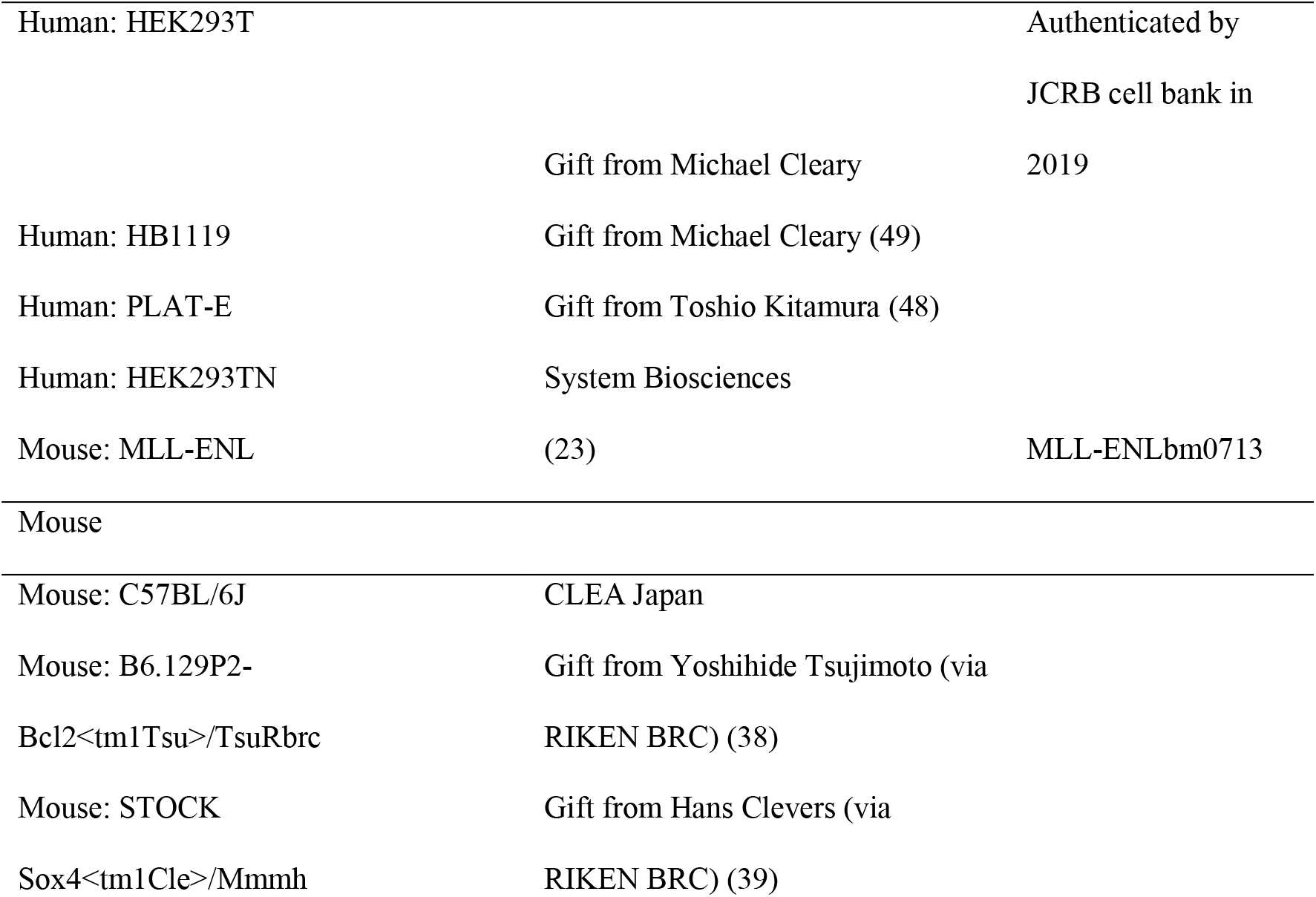
Reagents details

**Supplemental Table 2.**
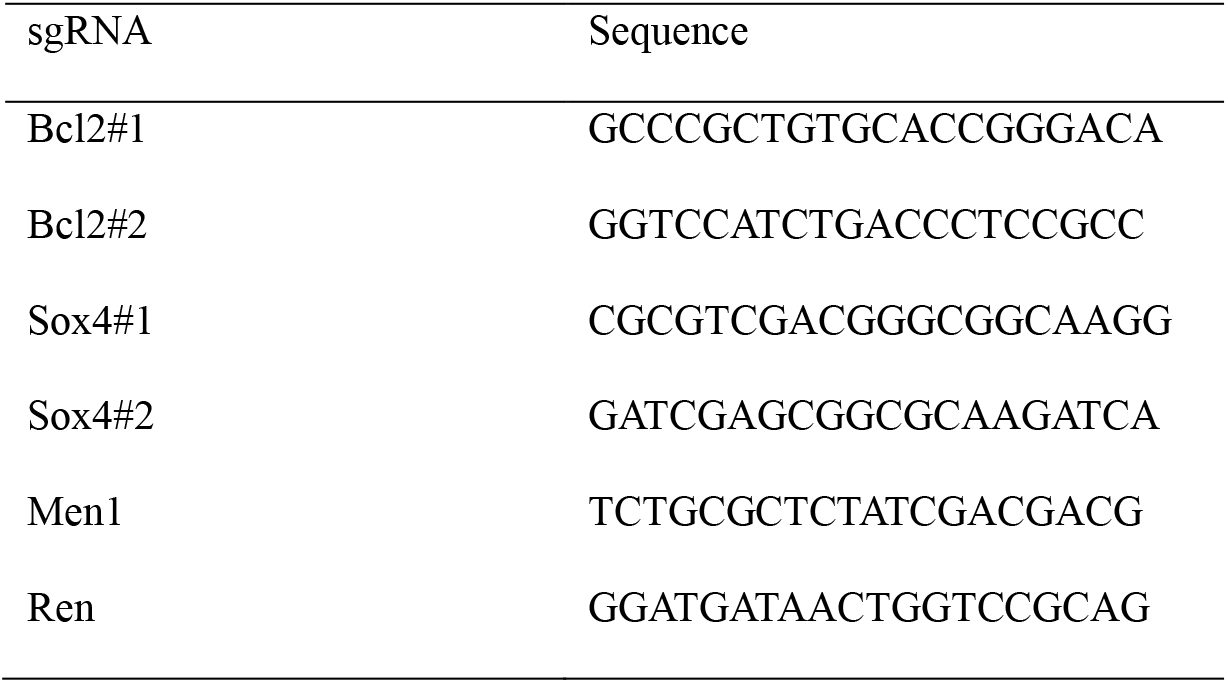
sgRNA sequences

**Supplemental Table 3.**
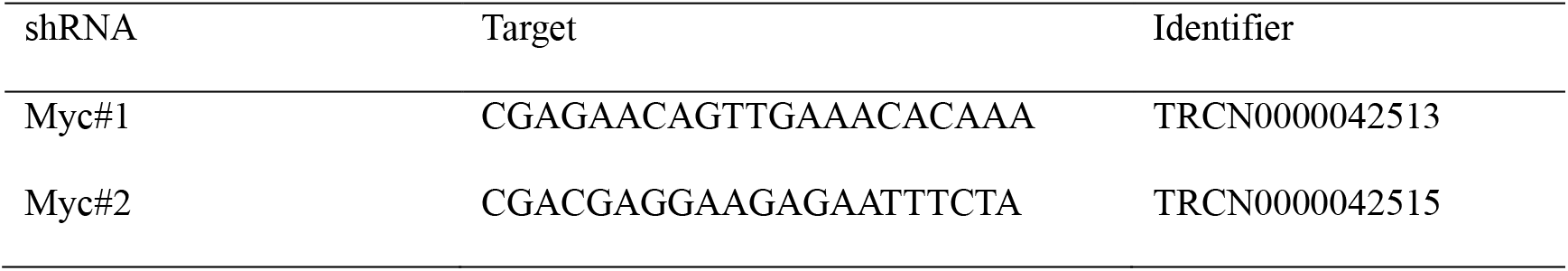
shRNA sequences

**Supplemental Table 4.**
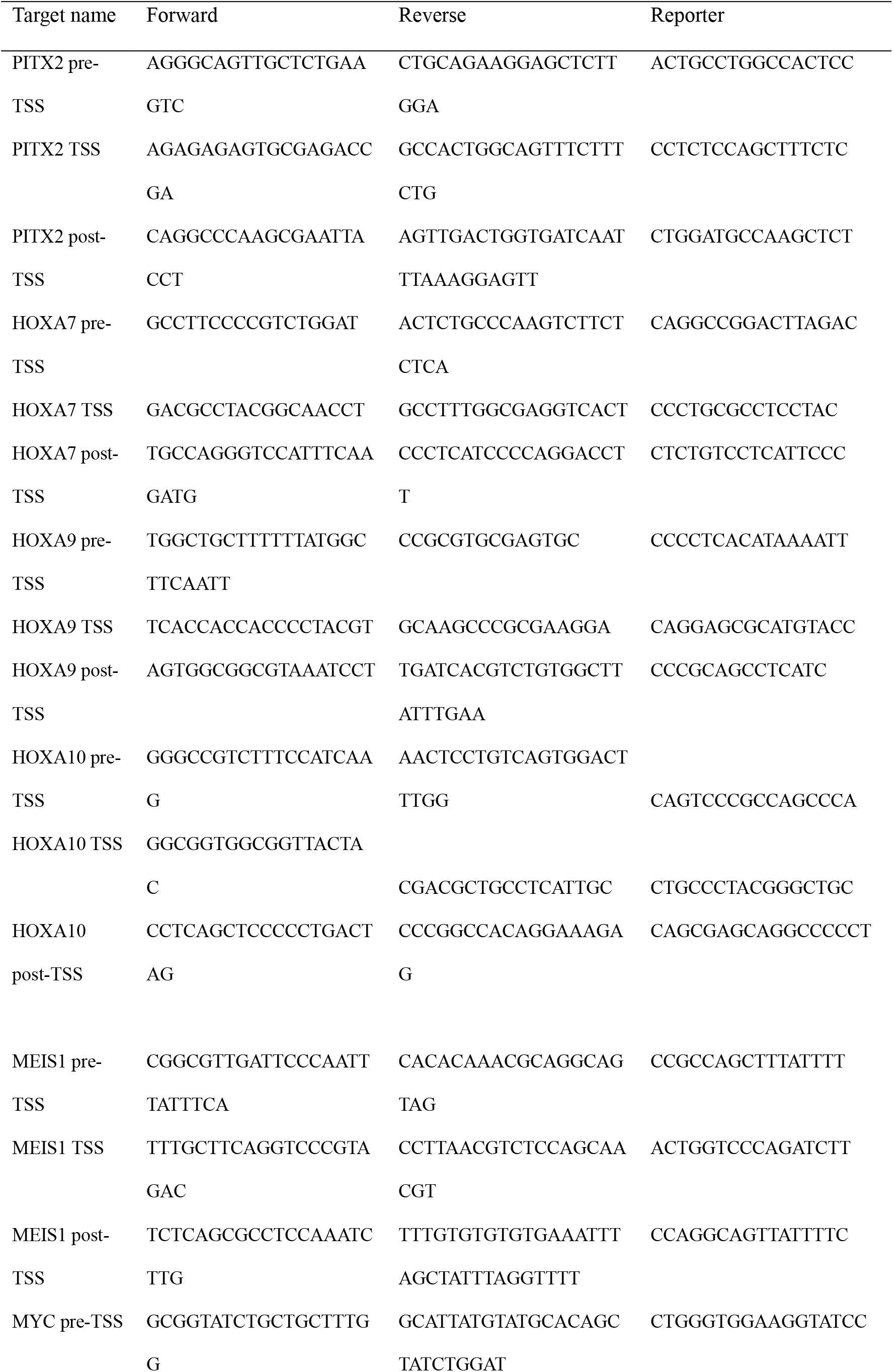
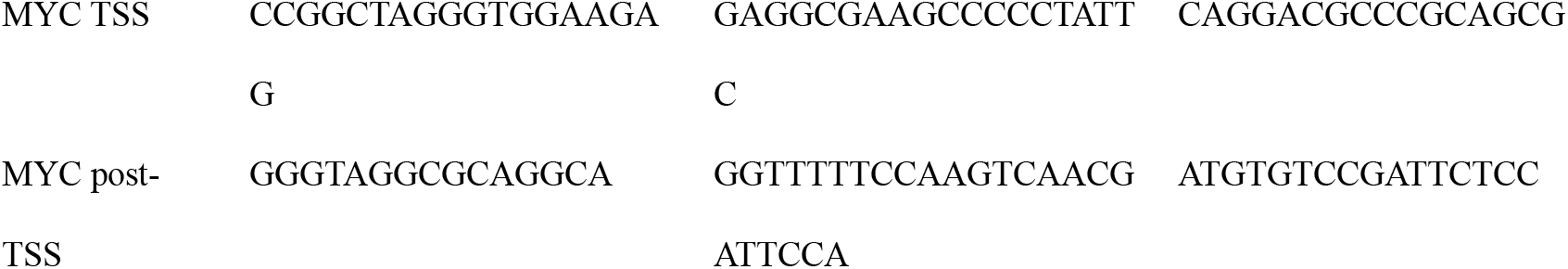
Custom qPCR probe/primer sequences

**Supplemental Table 5.**
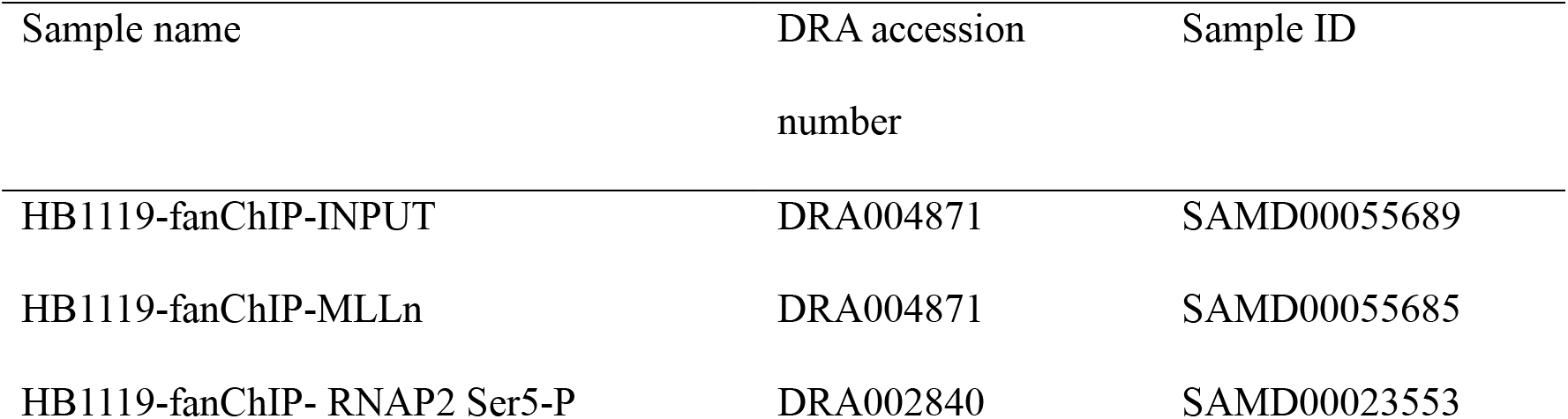
Accession numbers of ChIP-seq data

**Supplemental Table 6.** Accession numbers of mRNA-seq data

| Sample name | DRA accession number | Sample ID |
| --- | --- | --- |
| MP-fMLL-AF10-RNA 150916 | DRA010090 | SAMD00220959 |
| MP-fMLL-AF10-RNA 170719-1 | DRA010090 | SAMD00220960 |
| MP-fMLL-AF10-RNA 170719-2 | DRA010090 | SAMD00220961 |
| MP-fHOXA9-RNA 140824 | DRA010090 | SAMD00220962 |
| MP-fHOXA9-RNA 150812 | DRA010090 | SAMD00220963 |
| MP-fHOXA9-RNA 170719-1 | DRA010090 | SAMD00220964 |
| MP-fMYC-RNA 131204 | DRA010090 | SAMD00220965 |
| MP-fMYC-RNA 141113 | DRA010090 | SAMD00220966 |
| MP-fMYC-RNA 170719 | DRA010090 | SAMD00220967 |

